# Hyaluronidase-1-mediated glycocalyx impairment underlies endothelial abnormalities in polypoidal choroidal vasculopathy

**DOI:** 10.1101/2021.10.06.463357

**Authors:** Kan-Xing Wu, Natalie Jia Ying Yeo, Chun-Yi Ng, Florence Wen Jing Chioh, Fan Qiao, Xianfeng Tian, Yang Binxia, Gunaseelan Narayanan, Hui-Min Tay, Han-Wei Hou, N Ray Dunn, Xinyi Su, Chui Ming Gemmy Cheung, Christine Cheung

## Abstract

**Background:** Polypoidal choroidal vasculopathy (PCV), a subtype of age-related macular degeneration (AMD), is characterized by polyp-like dilatation of blood vessels and turbulent blood flow in the choroid of the eye. Gold standard anti-vascular endothelial growth factor (anti-VEGF) therapy often fails to regress polypoidal lesions in patients. Current animal models have also been hampered by their inability to recapitulate such vascular lesions. These underscore the need to identify VEGF-independent pathways in PCV pathogenesis.

**Results:** We cultivated blood outgrowth endothelial cells (BOECs) from PCV patients and normal controls to serve as our experimental disease models. When BOECs were exposed to heterogeneous flow, single-cell transcriptomic analysis revealed that PCV BOECs preferentially adopted migratory-angiogenic cell state, while normal BOECs undertook proinflammatory cell state. PCV BOECs also had a repressed protective response to flow stress by demonstrating lower mitochondrial functions. We uncovered that elevated hyaluronidase-1 in PCV BOECs led to increased degradation of hyaluronan, a major component of glycocalyx that interfaces between flow stress and vascular endothelium. Notably, knockdown of hyaluronidase-1 in PCV BOEC improved mechanosensitivity through activation of Krüppel-like factor 2, a flow-responsive transcription factor, which in turn modulated PCV BOEC migration. Barrier permeability due to glycocalyx impairment in PCV BOECs was also reversed by hyaluronidase-1 knockdown. Correspondingly, hyaluronidase-1 was detected in PCV patient vitreous humor and plasma samples.

**Conclusions:** Hyaluronidase-1 inhibition could be a potential therapeutic modality in preserving glycocalyx integrity and endothelial stability in ocular diseases with vascular origin.

## Background

Polypoidal choroidal vasculopathy (PCV), known to be a subtype of age-related macular degeneration (AMD), and is a major cause of vision loss in elderly populations. PCV is distinguished by the pathological presence of choroidal vessel networks with terminal polypoidal dilatations [1, 2]. The clinical diagnosis of PCV is confirmed by the visualization of vascular dilatation as hyperfluorescent nodules under indocyanine green angiography or orange-red subretinal nodules in routine ophthalmoscopic examinations [3–7]. Similar to typical neovascular AMD, PCV patients suffer from serosanguineous pigment epithelial detachment and submacular exudations resulting in a gradual loss of visual acuity [8, 9]. A background of choroidal vascular hyperpermeability is more frequently reported in PCV than in AMD [10]. Importantly, response of PCV to anti-vascular endothelial growth factor (anti-VEGF) therapy has been less consistent. In particular, while anti-VEGF controls the exudation, the underlying polypoidal lesion often fails to regress [8]. This current lack of effective therapeutic options for PCV that is refractory to anti-VEGF treatments reflects unresolved questions in the etiology of PCV.

Epidemiologic studies of PCV to date revealed a greater prevalence of PCV in populations of Asian (22-55% of neovascular AMD cases) and African descent [11] than Caucasian populations (8-13% of neovascular AMD cases) [12, 13]. Several cohort-based genome-wide association studies (GWAS) have revealed neovascular AMD to be a multifactorial disease with many single nucleotide polymorphisms (SNPs) identified to be significantly associated with the risk of disease development [14, 15]. However, GWAS studies have reported similar associations of SNPs in both PCV and neovascular AMD subtypes, with only rs10490924 in the age-related maculopathy susceptibility 2/high-temperature requirement A serine peptidase 1 (*ARMS2/HTRA1*) region showing significantly stronger association with PCV than neovascular AMD [14, 16–18]. The current understanding suggests that pathogenesis of PCV is likely an interplay of polygenic, biological and environmental factors [19].

The defining clinical feature of vascular dilatation in PCV suggests a unique perturbation to blood flow experienced by the choroidal endothelia. Indeed, optical coherence tomography angiography of PCV eyes, in combination with variable interscan time analysis revealed the varied nature of blood flow velocities within polyps with the center of polyps experiencing slower flow than the periphery, thus providing evidence for non-uniform flow within these vascular dilatations [20, 21]. The causes of these vascular malformations and the effects of non-uniform blood flow on the endothelium of these polyps are currently unknown.

Moreover, while several murine models for macular degeneration have been described, the mouse eye lacks a defined macula [22, 23] and laser-induced choroidal neovascularization models largely do not recapitulate polypoidal lesions in PCV [24]. In order to capture some of these genetic and environmental complexities in a human-relevant disease model, we leveraged on the use of patient-derived blood outgrowth endothelial cells (BOECs). BOECs can be derived *in vitro* from circulating endothelial colony-forming cells that originate from bone marrow or vessel resident stem cells [25–29]. With minimal manipulation, these cells give rise to mature endothelial cells in culture that are more likely to retain the genetic and epigenetic landscape of individuals [30]. Since it is well-known that endothelial cells are mechano-sensors that respond to shear stress by blood flow [31], we subject BOECs to variable and pulsatile flow conditions in order to recapitulate the dynamics within PCV polyps. Through single-cell analysis, we were able to discern transcriptional signatures of endothelial cells in response to heterogeneous flow and show that PCV and normal BOECs adopt distinct cell states under these conditions. Our findings demonstrate the powerful utility of patient-derived BOECs in modelling a complex vascular disease and illuminate molecular differences that can underlie the pathogenesis of PCV.

## Results

### Derivation and characterization of human blood outgrowth endothelial cells

We developed our BOEC models from peripheral blood mononuclear cell (PBMC) fractions isolated from PCV and normal donors according to established protocol [32]. Early colonies of BOECs emerged generally 7-14 days post-seeding of PBMCs (Fig. 1a). BOEC colonies were expanded for one week prior to passaging. On average for both normal and PCV groups, we obtained 1-3 colonies from every 10 million PBMCs (Fig. 1b). The proliferation capacity of BOECs were monitored from passages 3-8 when most of the BOEC lines demonstrated steady cell population doubling time (Fig. 1c). In our experimentations, we excluded potentially senescent BOECs if there was substantiate increase in their cell doubling time. To confirm endothelial identity, our derived BOECs were highly enriched for endothelial cell markers such as CD31 (> 99%) and CD144 (> 94%), but they had negligible expressions for leukocyte markers CD45 and CD68, and progenitor cell marker CD133, suggesting purity of our BOEC cultures (Fig. 1d). We further performed functional characterization of BOECs. PCV and normal BOECs were able to form tubular networks and showed comparable attributes (i.e. number of junctions, number of loops, branching length) with the positive control, human umbilical vein endothelial cells (HUVECs) (Fig. 1e). In a three-dimensional fibrin gel bead sprouting assay, our BOECs also displayed sprouting with filopodia, characteristic of endothelial protrusions in mediating guidance cues during angiogenesis (Fig. 1f).

**Figure 1:**
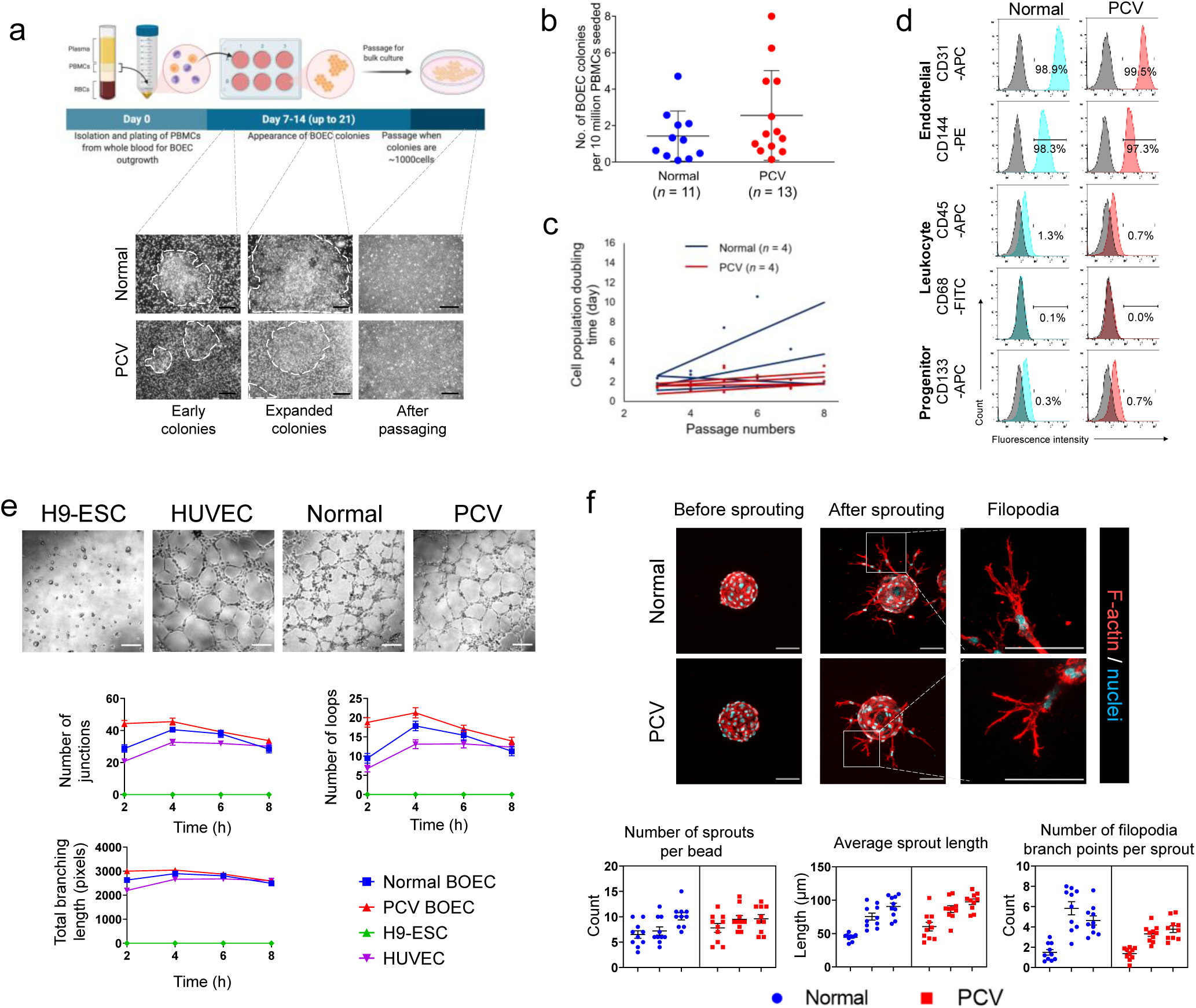
Derivation and characterization of human blood outgrowth endothelial cells. **(a)** Workflow illustrating the generation of BOECs. Images show BOEC colonies emerging during days 7-14 post-seeding of PBMCs, followed by characteristic cobblestone-like endothelial cells after passaging of colonies (scale bar, 100 µm). **(b)** Number of BOEC colonies per million PBMCs obtained from normal controls and PCV patients. **(c)** Proliferation dynamics of BOEC lines measured by cell doubling duration over passages. **(d)** Flow cytometry characterization of BOECs for endothelial, leukocyte and progenitor cell markers (grey - isotype control; red/blue – cell lineage marker staining). **(e)** Tube formation assay with representative images of tube formation ability of BOECs at 4h. Bottom panel shows quantification of junctions, loops (tubes) and total branching length (total length of loops and branches) over time (quantified from *n*=12 optical fields per timepoint from each cell line). H9-stem cells (H9-ESC) and HUVECs are negative and positive controls for tube formation, respectively. Data are mean ± s.e.m. Scale bars, 200 µm. **(f)** Fibrin gel bead sprouting assay of BOECs at 24h. BOECs were immunostained for F-actin (red) and DAPI (cyan). Bottom panel shows measurements of relevant sprouting parameters (quantified from *n*=10 beads from each individual, BOECs from 3 PCV and 3 normal individuals). Data are mean ± s.e.m. Scale bars, 100 µm.

Based on marker expressions and functional characterization, both PCV and normal BOECs demonstrated comparable attributes. Extrinsic factors such as complement dysregulation and oxidative stress, as well as retinal pigment epithelial (RPE) cells being one of the main producers of VEGF in the eye, play key roles in pathological endothelial behaviors [33]. To understand RPE cells’ paracrine effects on endothelial function, induced pluripotent stem cells (iPSCs) were created through Sendai-based reprogramming of PBMC samples from PCV and normal individuals. We generated 4 iPSC lines from 2 PCV patients and 4 iPSC lines from 2 normal individuals (Supplemental Table S1). All PCV lines were homozygous for the risk alleles in *ARMS2/HTRA1* locus, and heterozygous at CFH locus. All the normal lines harboured protective alleles in at least 1 locus. The derived iPSC lines showed tightly packed colonies and expressed classical markers of pluripotency, as well as germ layer markers upon embryoid body differentiation (Supplemental Figure S1a). RPE differentiation was then performed on PCV and normal iPSCs based on established protocols [34] to derive iPSC-RPE cells which developed pigmentation from day 34 of differentiation onwards (Supplemental Figure S1b). After monolayers of iPSC-RPE cells were re-seeded onto transwell to induce polarity, we confirmed their barrier integrity, typical of RPE cells, by trans-epithelial electrical resistance measurements over time until day 71 post-seeding (Supplemental Figure S1c). To validate RPE cells’ contribution to secreted VEGF, we found that iPSC-RPE conditioned media from the basal compartments had seemingly greater amount of VEGF than that from the apical compartments (Supplemental Figure S1d), consistent with RPE cell biology. However, we did not observe difference in VEGF production between PCV and normal iPSC-RPE cells based on the current number of cell lines. Likewise, there was no significant difference in VEGF levels between PCV and normal plasma samples (Supplemental Figure S1e). We postulated that there could be other systemic mediators affecting endothelial health. Hence, we exposed PCV and normal BOECs to autologous plasma and found that some angiogenic attributes could be intensified by plasma stimulation (Supplemental Figure S1f). As there is a multitude of paracrine influences which are known to cause endothelial dysfunction, the knowledge gaps in PCV endothelial cell autonomous effects remain understudied. We hereby focused on deciphering intrinsic endothelial mechanisms which could be further interrogated in our PCV BOEC disease model.

### Blood outgrowth endothelial cells adopt diverse cell states under heterogeneous flow

Among our derived BOEC lines (13 PCV, 11 normal), we prioritized those that passed quality controls in terms of endothelial marker expressions and functional attributes (Fig. 1). In addition, we genotyped the BOEC lines for AMD/PCV genetic risk loci in *ARMS2/HTRA1* (rs10490924 and rs11200638) and *CFH* (rs800292). Collectively, we selected 4 PCV and 6 normal lines for further experimentation (Supplemental Table S1). Three PCV donors were homozygous for the risk alleles in *ARMS2/HTRA1* locus, and 2 PCV donors were heterozygous at *CFH* locus. All the normal controls harboured protective alleles in at least 1 locus.

We introduced heterogeneous flow as a stress paradigm to PCV and normal BOECs. To recapitulate the variable flow conditions in PCV polyps, we utilized an orbital flow setup to generate a continuum of shear forces with high magnitude, uniaxial shear stresses at the edge of wells and low magnitude, multidirectional shear stresses in the center [35]. PCV and normal BOECs were exposed to 24h of rotation (Fig. 2a). Peak fluid shear in our setup was estimated to be around 10 dyne/cm^2^ using 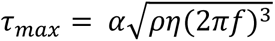, with α being the orbital radius (0.1cm), ρ as density of medium (assumed 0.9973g/mL) [36], η as the medium viscosity (assumed 0.0101 poise) [36] and *f* as the frequency of rotation (210/60rps) [36, 37]. In a similar setup using a 6-well plate, optical Doppler velocimetry measured shear stress of 5 dyne/cm^2^ at the center of the well and 11 dyne/cm^2^ at the periphery [36]. This range of shear stress magnitudes is well within reported physiological range where shear stresses have been described at ±4 dyne/cm^2^ around curvatures, bifurcations and branches, while straight arterial regions experience shear forces of approximately 10 – 20 dyne/cm^2^, at times reaching 40 dyne/cm^2^ [38]. The effects of this heterogeneous flow can be seen from the staining of vascular endothelial cadherin (CDH5) (Fig. 2b), where both PCV and normal BOECs demonstrated alignment to flow direction at well periphery but not in the center. Correspondingly, caveolin-1 (CAV1) which forms part of the endothelial mechanosensing machinery [39], showed re-distribution to BOEC cell edges as a response to higher shear stress at the well periphery (Fig. 2b).

**Figure 2:**
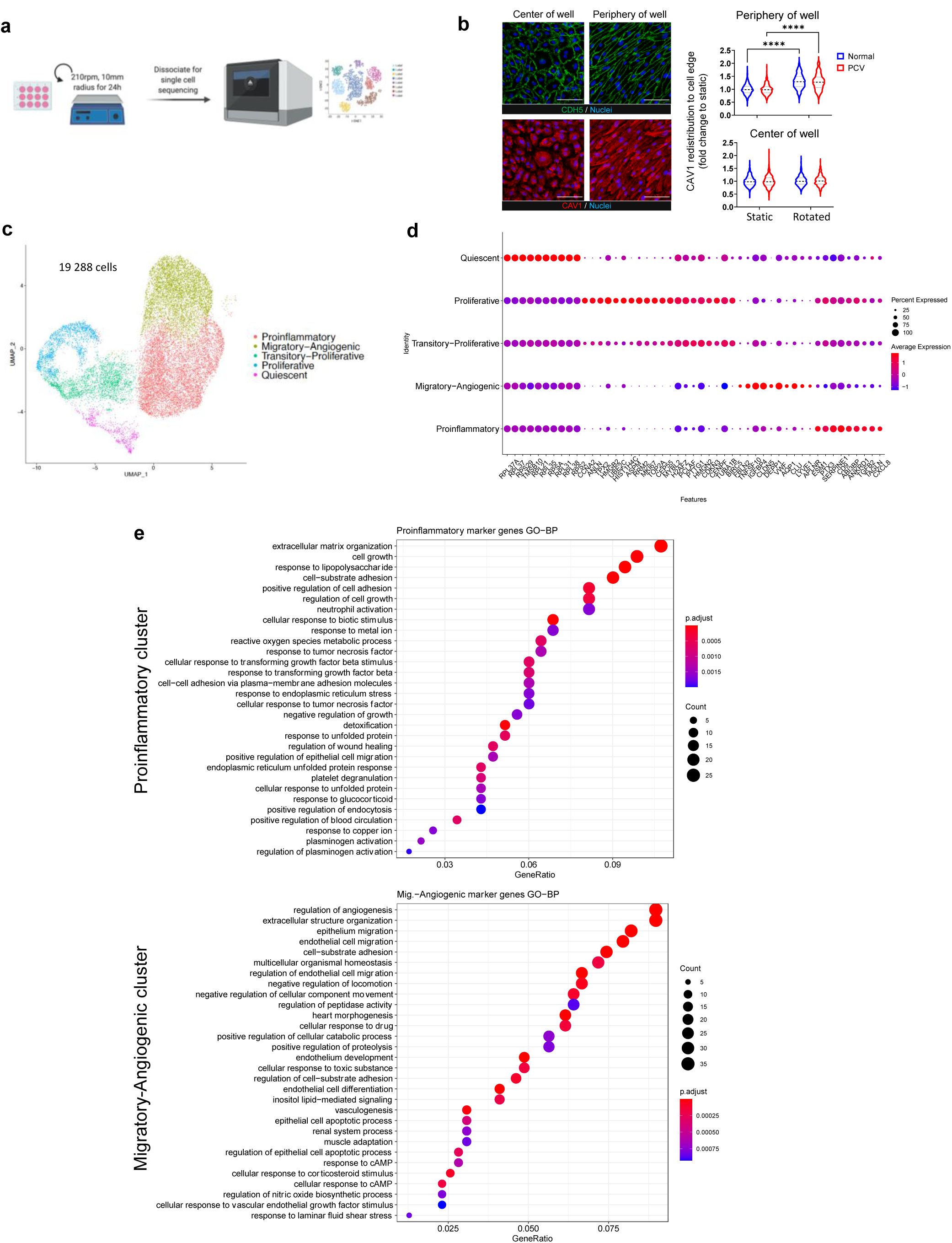
Single-cell RNA sequencing reveals different cell states after exposure to heterogeneous flow. **(a)** Workflow showing orbital flow setup used on PCV and normal BOEC lines prior to dissociation for scRNA sequencing. **(b)** CDH5 (green) and CAV1 (red) immunostainings reveal patterns of heterogeneous flow in the setup. Nuclei were stained with DAPI (blue). Right panel shows measurements of CAV1 intracellular re-distribution as an indicator of flow response (BOECs from 4 PCV and 3 normal individuals). Data shown are normalized to individual static conditions with median and quartiles indicated. *p* values were obtained using two-way ANOVA with Tukey’s multiple comparisons test. \*\*\*\**p*<0.0001. Scale bar, 100 µm. **(c)** Single-cell UMAP showing sequenced cells for all subjects (*n*= 19,288 cells, BOECs from 2 PCV and 2 normal individuals). We identified 5 distinct transcriptomic cell states across the integrated dataset, 1) Proinflammatory, 2) Migratory-Angiogenic, 3) Transitory-Proliferative, 4) Proliferative and 5) Quiescent cells. **(d)** Gene expression of top 10 positive markers genes for each of the cell states shown in **(c)**. **(e)** Gene set enrichment analysis against GO (Biological processes) database of marker genes for two largest clusters shown here ranked by Bonferroni corrected *p*-values.

To uncover the molecular underpinning of PCV endothelial abnormalities, we performed single-cell RNA sequencing (scRNA-seq) to resolve transcriptomic differences between PCV and normal BOECs in response to heterogeneous flow. Our single-cell analysis revealed 5 clusters of transcriptionally distinct cell states that we classified as 1) Proinflammatory, 2) Migratory-Angiogenic, 3) Transitory-Proliferative, 4) Proliferative and 5) Quiescent cells (Fig. 2c). These cell states were determined firstly by analysis of cluster-enriched marker genes. For example, representative genes such as inflammatory markers *CXCL8* and *TGFβ2*, as well as cell cycle/ proliferation markers *MKI67*, *PCLAF* and *TOP2A* were among the top 10 cluster-enriched marker genes where the highest average expression for each gene corresponded to their identified cell states (Fig. 2d). Furthermore, expression patterns of cell state-defining marker genes were conserved between PCV and normal datasets (Supplemental Fig. S2c).

Secondly, we confirmed cell state identities by the predominant processes found from gene enrichment analyses of top cluster-specific marker genes using Gene Ontology [40, 41] and Reactome [42] databases. Enriched processes for the Proinflammatory cell state included myeloid leukocyte adhesion, neutrophil activation and platelet degranulation, while those for Migratory-Angiogenic cell state included positive regulation of endothelial cell migration, angiogenesis and hyaluronan (HA) uptake and degradation (Fig. 2e, Supplemental Fig. S2d and e). The Transitory-Proliferative and Proliferative cell states shared several processes such as nuclear division and cell cycle (Supplemental Fig. S2d and e). Taken together, both PCV and normal BOECs adopted diverse transcriptomic cell states in response to heterogeneous flow.

### PCV and normal endothelial cells demonstrate differential responses to heterogeneous flow

In discerning meaningful differences between PCV and normal BOECs, we found a majority of normal BOECs adopt Proinflammatory cell state (50.36%) under flow treatment. Notably, we noticed a distinct departure in PCV BOECs, with the majority of cells found in Migratory-Angiogenic cell state instead (41.69%) (Fig. 3a). These differences represented a primary shift between Proinflammatory and Migratory-Angiogenic cell states as the other cell states remain comparable between PCV and normal in terms of cell proportions. These changes in cell proportions were found largely conserved across each of the PCV and normal BOEC samples, ruling out bias arising from inter-individual variabilities (Supplemental Fig. S3a).

**Figure 3:**
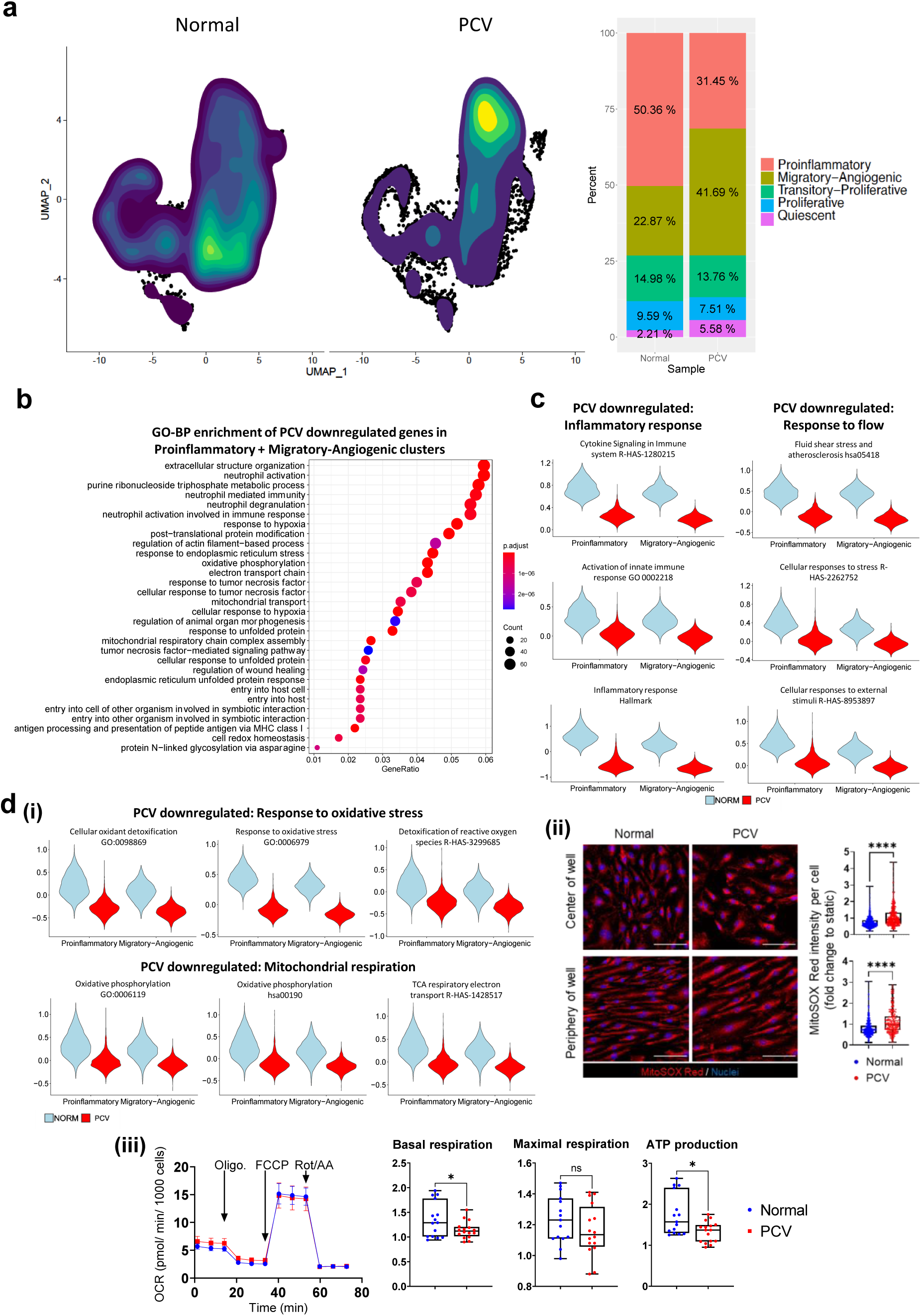
PCV endothelial cells show attenuated response to heterogeneous flow. **(a)** Contour plot overlays on UMAPs of normal and PCV samples showing the density distribution of sequenced cells across all clusters. Right panel shows the percentage breakdown of cells per cell state for normal and PCV samples. **(b)** Differential expression analysis was carried out comparing PCV and normal cells in the Proinflammatory and Migratory-angiogenic cell states. Gene set enrichment analysis against GO (Biological processes) database reveals enriched processes from PCV downregulated genes shown here ranked by Bonferroni corrected *p*-values. **(c)** Gene set enrichment analyses of PCV downregulated genes in these clusters were expanded to include Reactome, KEGG and MSigDB databases. Significantly enriched pathways or processes (adj. *p*<0.01) are shown here as violin plots that represent calculated module scores for the gene sets indicated. **(d)** (**i**) Module scores of PCV and normal cells for significantly enriched oxidative stress response and mitochondrial respiration gene sets (adj. *p*<0.01) are shown as violin plots here. (**ii**) MitoSOX Red staining of PCV and normal BOECs after 24h of heterogeneous flow. Nuclei were stained DAPI (blue). Right panel shows intensity per cell measurements of MitoSOX Red staining (*n* > 1,000 cells anaylzed per group, BOECs from 2 PCV and 2 normal individuals). Data shown are normalized to individual static conditions with median and quartiles indicated. *P* values were from two-tailed Mann Whitney test . \*\*\*\**p*<0.0001. Scale bar, 100 µm. (**iii**) Mitochondrial function assay of BOECs after 24h of heterogeneous flow. Left panel shows oxygen consumption rate (OCR) of BOECs in response to oligomycin (Oligo.), carbonyl cyanide-4 (trifluoromethoxy) phenylhydrazone (FCCP) and rotenone/ antimycin A (Rot/AA). Right panel shows measurements per well of basal respiration, maximal respiration and mitochondrial ATP production (BOECs from 4 PCV and 3 normal individuals). Data shown are normalized to individual static conditions with median and quartiles indicated. *P* values for basal respiration and maximal respiration were from two-tailed *t* tests with Welch’s correction and for ATP production from two-tailed Mann-Whitney test, \**p*<0.05.

Next, we performed differential expression analysis between PCV and normal datasets with a focus on combined Proinflammatory and Migratory-Angiogenic cell states. Gene set enrichment analyses of PCV downregulated genes revealed largely inflammation-related events such as neutrophil activation and neutrophil-mediated immunity (adj. *p*< 0.01) (Fig. 3b), indicating that PCV BOECs expressed a weaker proinflammatory profile than their normal counterparts. Gene set enrichment analysis was expanded to include databases from KEGG [43] and MsigDB [44] to reveal enrichment of key pathways such as Cytokine Signaling in Immune system (R-HAS-1280215, Reactome), Inflammatory response (Hallmark, MSigDB), Fluid shear stress and atherosclerosis (hsa05418, KEGG) and Cellular responses to external stimuli (R-HAS-8953897). Module scores for the average expression of genes in these pathways demonstrated the degree of differential expression between PCV and normal BOECs for these gene sets (Fig. 3c). Upregulation of proinflammatory genes (e.g. *CXCL8*, *ICAM*) are well-described processes in endothelial cells subjected to disturbed, multidirectional flow [45]. Our findings suggested reduced sensitivity to flow by PCV BOECs in contrast to normal BOECs.

In addition, we found significant enrichment in oxidative stress response and oxidative phosphorylation related processes with PCV BOECs expressing lower levels of genes involved in detoxification of reactive oxygen species and respiratory electron transport chain Fig. 3d(i). MitoSOX Red staining for superoxide and mitochondrial activity measurements validated these transcriptomic findings with PCV BOECs showing significantly higher levels of oxidative stress and lower mitochondrial functions than normal BOECs (Fig. 3d(ii) and (iii)) (*p*<0.05). Antioxidant response genes such as superoxide dismutases (*SOD1*) are upregulated in endothelial cells as a protective response to oxidative stress in oscillatory flow [45]. Our MitoSOX staining and lower mitochondrial functions might explain a repressed protective response in PCV BOECs towards heterogeneous flow.

### PCV endothelial cells have increased migratory capacity and barrier permeability

In addition to reduced flow response, PCV BOECs had a stronger migratory transcriptomic profile than normal BOECs. Our gene set enrichment analyses of the PCV upregulated genes showed that the major enriched processes revolved around cell migration and cell locomotion, in particular blood vessel endothelial cell migration (GO:0043534) (Fig. 4a). To functionally validate these transcriptomic differences, we went on to assess the migratory capacity of PCV and normal BOECs in a wound healing assay. PCV BOECs demonstrated significantly greater wound closure than normal BOECs with and without exposure to orbital flow, although the differences between PCV and normal were greater after 24h of heterogeneous flow (23.17% ± 3.892, *p*<0.0001) than static cultures (20.75% ± 5.865, *p*=0.0046) (Fig. 4b). Intimately linked to the process of cell migration, extracellular matrix (ECM) modifying processes were also found to be significantly enriched in the PCV upregulated gene set (Fig. 4c), in particular proteases such as *MMP1*, *MMP16* and *ADAMTS18* (Supplemental Fig. 3b).

**Figure 4:**
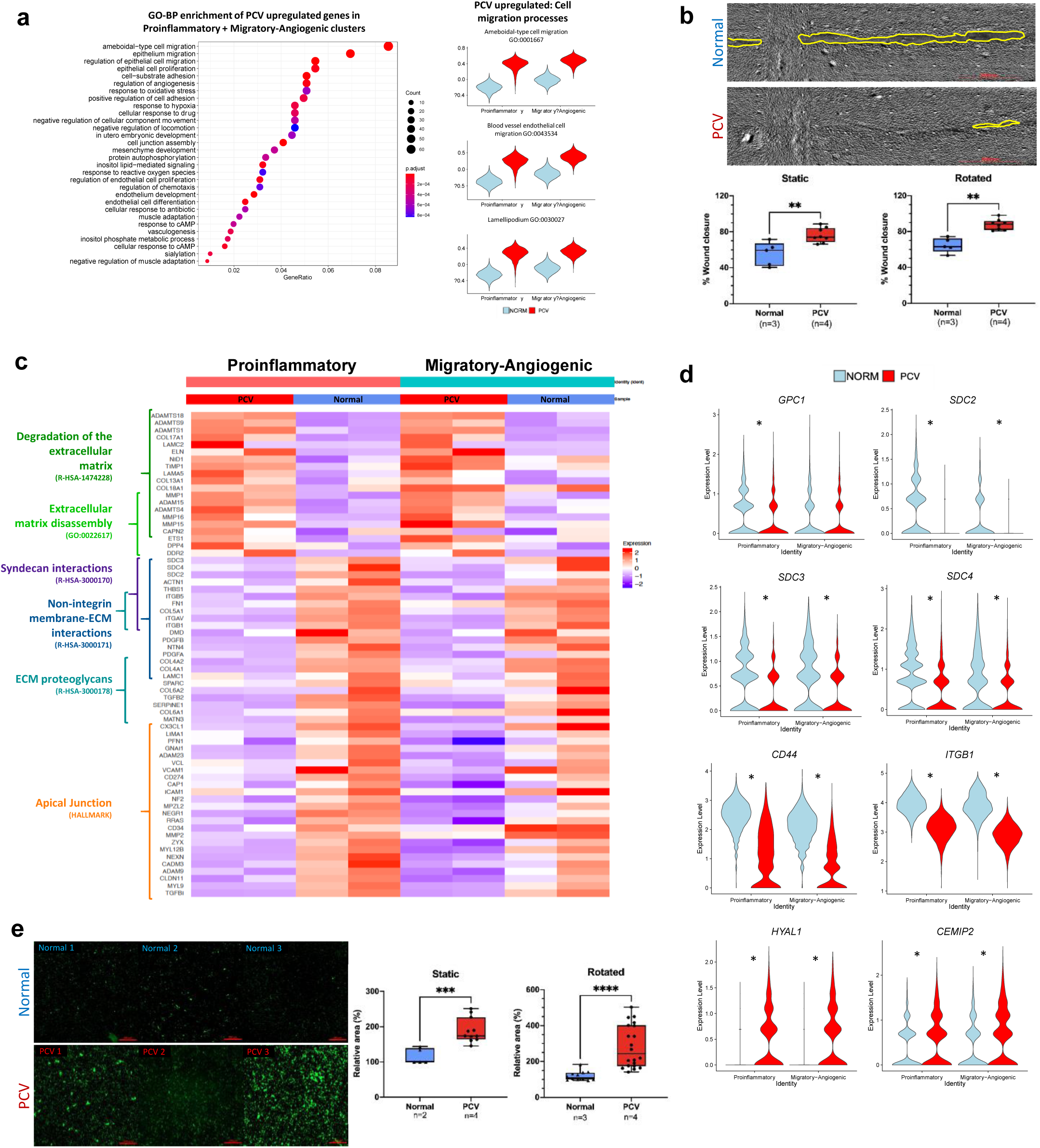
Phenotypic characterization of migratory capacity and barrier permeability in PCV endothelial cells. **(a)** Gene set enrichment analysis against Gene Ontology (Biological processes) database reveals enriched processes from PCV upregulated genes shown here ranked by Bonferroni corrected *p*-values. Right panel shows violin plots that represent calculated module scores for the processes indicated. **(b)** Wound healing assays carried out to assess migratory capacity of BOECs 21h post-scratch after 24h of static or rotated (heterogeneous flow) culture. Left panel shows representative images from heterogeneous flow condition with yellow outlines indicating cell-free wound regions after 21h, scale bar, 2000µm. Box and whiskers plots showing median with minimum and maximum. *n* indicates number of cell lines evaluated for each group while *p*-value is from two-tailed t-test, ** *p*<0.01. **(c)** Heatmap showing average expression of cells from each individual BOEC sample in the indicated cell states. **(d)** Violin plots of glycocalyx-related genes found to be significantly differentially expressed between PCV and normal BOECs, *adj. *p*< 0.01. **(e)** *In vitro* vascular permeability imaging assay using biotinylated-gelatin coated surfaces. Intracellular gaps were revealed by Neutravidin-FITC staining after 24h of static or rotated (heterogeneous flow) culture. Shown are representative frames from different BOEC lines, scale bar, 200µm. Box and whiskers plots showing median with minimum and maximum. Area was normalized against average FITC area of normal BOEC lines. *n* indicates number of cell lines evaluated for each group while *p*-value is from two-tailed t-test, *** *p*<0.001, **** *p*<0.0001.

In linking reduced flow response to increased migratory capacity in PCV BOECs, we hypothesized a perturbed extracellular milieu that interfaced between flow and endothelial cells. Further to the enrichment of ECM-degrading processes in PCV upregulated genes, syndecan interactions (R-HAS-3000170) was found to be significantly downregulated in PCV BOECs (Fig. 4c). Syndecans are part of the endothelial glycocalyx, which is a mechanosensing meshwork of glycosaminoglycans covering the luminal surface of endothelial cells, held covalently by proteoglycan core proteins^40^. Its apical positioning and extensive coverage of the cell surface enables sensing and transduction of hemodynamic forces to interacting partners on the plasma membrane [46, 47]. Here, we looked at the expression profiles of glycocalyx-related genes and found significantly lowered expression in PCV for heparan sulfate (HS) core proteins (*GPC1*, *SDC2*, *SDC3* and *SDC4*) and HA core protein (*CD44*), while genes for HA degrading enzymes *(HYAL1* and CEMIP2) were significantly higher in PCV (Fig. 4d). Also, β1 integrin (*ITGB1*), a reported mechano-sensor of blood flow usually upregulated in the presence of flow [48] was found to be downregulated in PCV relative to normal BOECs.

Phenotypically, we used an image-based permeability assay and demonstrated significantly larger areas of intracellular gaps in PCV BOEC monolayers than normal BOEC monolayers in both static and rotated conditions (Fig. 4e). Similar to the wound healing migration assay, differences between PCV and normal BOECs in barrier permeability became exacerbated after heterogeneous flow treatment (163.0% ± 34.91, *p*<0.0001) than without (74.09%± 17.02, *p*=0.0007). These results revealed fundamental differences between PCV and normal BOECs in endothelial functions in the forms of barrier integrity and migratory capacity, which were amplified by disturbed flow conditions.

### Increased HYAL1 levels in PCV endothelial cells impair glycocalyx

The integrity and composition of endothelial glycocalyx can determine the efficiency and extent of mechano-sensitivity and force transduction [46, 47]. Of the 6 known genes coding for hyaluronidases in humans [49], we found *HYAL1* to be expressed at a significantly higher level in PCV than normal BOECs across both Proinflammatory and Migratory-Angiogenic clusters (Fig. 4d and Supplemental Fig. S3b). Hence, we selected *HYAL1* for further validation due to the strong differential expression in the transcriptomic data and its active functional role in modifying glycocalyx composition through its hyaluronan (HA) degrading activities. Western blot analyses of BOEC lysates, subjected to heterogeneous flow, validated the increased expression of HYAL1 in PCV BOECs at the proteomic level (Fig. 5a).

**Figure 5:**
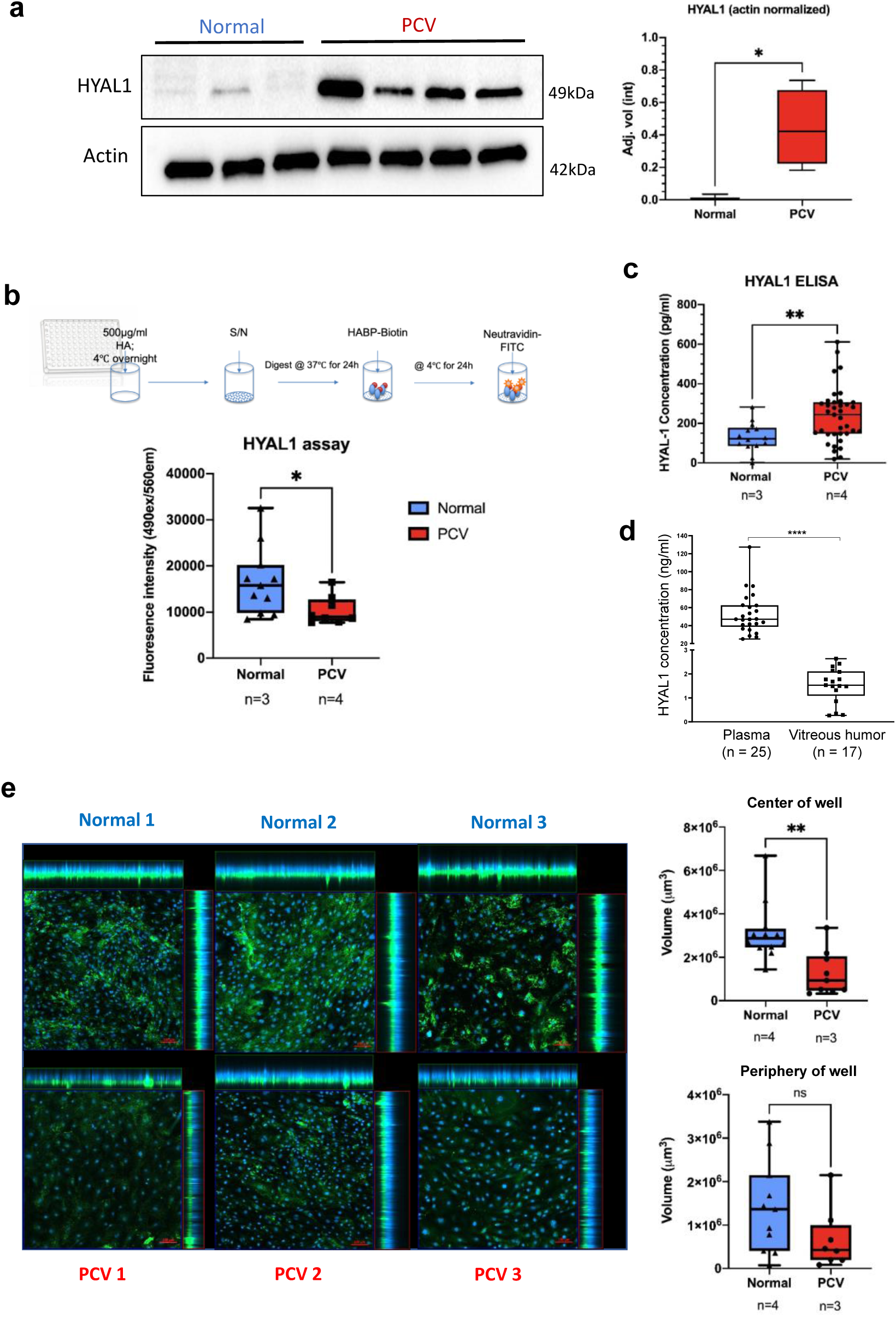
HYAL1 enzymatic activities and perturbation of glycocalyx hyaluronan in PCV endothelial cells. **(a)** Western blot analyses of cell lysates from 4 PCV and 3 normal BOEC lines, subjected to 24h of heterogeneous flow, showing HYAL1 detection at 51kDa and Actin as the loading control at 42kDa. Right panel shows densitomeric quantification of HYAL1 bands normalized against Actin. Box and whiskers plots showing median with minimum and maximum. *p*-value is from two-tailed t test, * *p*<0.05. **(b)** Enzymatic activity of HYAL1 was evaluated according to the workflow illustrated in the upper panel. Conditioned media (S/N, supernatant) was obtained from BOEC cultures rotated for 24h. Box and whiskers plots showing median with minimum and maximum. *n* indicates number of cell lines evaluated for each group while *p*-value is from two-tailed t-test, * *p*<0.05. **(c)** ELISA was used to detect HYAL1 in BOEC conditioned media after 24h heterogeneous flow. Box and whiskers plot shows median with minimum and maximum. *n* indicates number of cell lines evaluated for each group while *p*-value from two-tailed Mann-Whitney test, ** *p*<0.01. **(d)** ELISA detection of HYAL1 in PCV patient plasma and vitreous humor. Box and whiskers plot shows median with minimum and maximum. Every point represents an average of duplicates or triplicates of an individual patient. *n* indicates number of patients evaluated for each group while *p*-value from two-tailed Mann-Whitney test, **** *p*<0.0001. **(e)** HA in the glycocalyx was detected using biotinylated HABP before detection with FITC-Neutravidin (green). Nuclei were stained blue with DAPI. Left panel shows maximal intensity projections of representative frames from each cell line evaluated. Cell were subjected to heterogeneous flow for 24h before cold methanol fixation and staining. Three image frames were taken for each well region. Right panel shows total HA-stained areas summed across z-stacks per image frame. Stacks were taken to the point of last visible FITC signal in each frame. Box and whiskers plot showing median with minimum and maximum. *n* indicates number of cell lines evaluated for each group while *p*-value is from two-tailed t test, ** *p*<0.01.

HYAL1 is a secreted protein that is endocytosed and activated at low pH in lysosomes [50, 51]. As such, we probed the proteolytic activity of secreted HYAL1 and found greater degradation of HA (*p*=0.0194) using conditioned media from PCV BOECs subjected to heterogeneous flow (Fig. 5b). Higher levels of HYAL1 were also detected in these conditioned media of PCV BOECs as quantified by ELISA (Fig. 5c). Subsequently, HYAL1 was detectable in PCV patients’ plasma and eye vitreous humor extracts (Fig. 5d).

Using a biotinylated-HA binding protein (HABP), we were able to visualize the HA component of BOEC glycocalyx and observed an overall decrease in HA staining in PCV BOECs relative to normal BOECs after heterogeneous flow (Fig. 5e). Volumetric analyses of HA-staining across z-stacks revealed a significantly lower HA volume in PCV BOECs exposed to heterogeneous flow at the center of well (Fig. 5e). While we observed the same trend in BOECs at the periphery of well, significant difference of HA content between PCV and normal BOECs was not achieved. As aforementioned, flow conditions in the center of well had relatively lower shear stress and higher multi-dimensionality than that found at the periphery of well. Collectively, these results validated the transcriptomic data of higher *HYAL1* expression in PCV BOECs and suggest that PCV BOECs may experience a higher HA turnover and breakdown under pathological flow conditions.

### Modulation of HYAL1 restores normal cell migration and barrier integrity in PCV endothelial cells

Finally, we evaluated if the increased expression of HYAL1 in PCV BOECs can play a role in mediating the functional phenotypes of increased migratory capacity and barrier permeability. We used small-interfering RNA (siRNA) to silence gene expressions of *HYAL1* that were confirmed at protein levels in the human BOECs (Fig. 6a). HYAL1 knockdown (siHYAL1) was able to reduce wound closure percentage in PCV samples significantly (*p=*0.046) at 50nM, while normal BOECs remained unperturbed (Fig. 6b). The knockdown of HYAL1 was also able to restore PCV migratory capacity to the similar level as normal BOECs (Fig. 6b). We postulated that knockdown of HYAL1 might improve mechanosensing ability in PCV BOECs, in part through preserving HA in endothelial glycocalyx. Hence, we examined the flow-responsive transcription factor, Krüppel-like factor 2 (*KLF2*)[52], and found that *KLF2* expressions were indeed activated in BOECs exposed to heterogeneous flow compared to static condition (Fig. 6c). HYAL1 knockdown further upregulated *KLF2* level significantly in PCV BOECs under flow, but not in normal BOECs (Fig. 6c). This might explain the aforementioned observation that PCV BOEC migratory capacity could be effectively modulated by HYAL1 knockdown, possibly through activation of *KLF2*, which in turn exerted anti-migratory effect on endothelial cells[53]. Furthermore, HYAL1 knockdown significantly reduced barrier permeability in PCV BOECs to a level similar to normal BOECs transfected with non-targeting siRNA (NT) (*p*=0.0483) (Fig. 6d). Hence, HYAL1 modulation could reverse abnormal PCV endothelial cell migration and barrier permeability.

**Figure 6:**
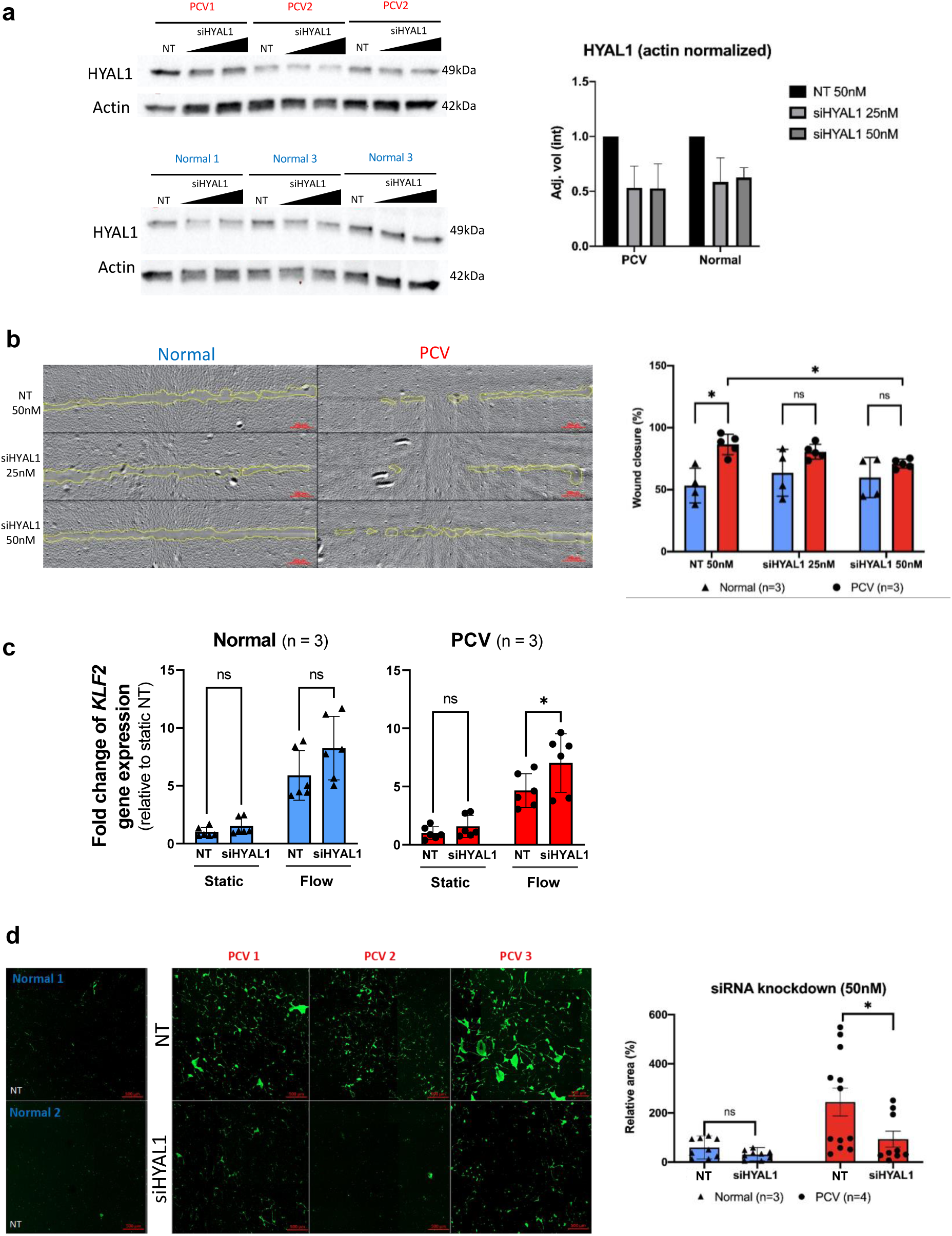
Modulation of HYAL1 normalizes abnormal cell migration and barrier permeability of PCV endothelial cells. **(a)** siRNA knockdown of *HYAL1* in BOECs. BOECs were transfected with siRNA for 4 days, including 24h heterogeneous flow treatment. Cell lysates were harvested and analyzed for HYAL1 levels. Bar graphs show mean actin-normalized HYAL1 band intensities with error bars representing standard deviations (siHYAL1, short-interfering RNA of *HYAL1*; NT, non-targeting siRNA). **(b)** Wound healing assays were carried out to assess migratory capacity of BOECs at 21h post-scratch after 24h of static or rotated culture. siRNA knockdown was carried out 48h prior to orbital rotation. Top panel shows representative image set from the flow condition with yellow outlines indicating cell-free wound regions at 21h post-scratch, scale bar, 200µm. **(c)** Relative *KLF2* gene expressions in BOECs in static condition and after 6 hours of heterogeneous flow exposure, with treatment of either NT or siHYAL1. **(d)** *In vitro* vascular permeability imaging assay using biotinylated-gelatin coated surfaces. Intercellular gaps were revealed by Neutravidin-FITC staining after 24h of static or rotated culture. siRNA knockdown was carried out 48h prior to orbital rotation. Shown are representative frames from different BOEC lines, scale bar, 500µm. Area was normalized against average FITC area of normal BOEC lines. All bar graphs showing means with standard deviations. *n* indicates number of cell lines evaluated for each group while *p*-value is from two-tailed t-test, \**p*<0.05.

## Discussion

We have addressed a major knowledge gap in the endothelial underpinning of PCV as most ocular disease modeling studies have focused on the biology of RPE cells. Previously, the difficulty of culturing human primary choroidal endothelial cells in comparison to human RPE cells could underlie the dearth of studies on the role of endothelial cells. Here, derivation of BOECs represents a minimally-invasive method of establishing disease-relevant endothelial cells for experimentations. We report single-cell analysis of the differential responses of human endothelial cells from healthy controls and PCV patients to heterogeneous flow. We found that PCV BOECs are abnormally migratory and have increased barrier permeability. This is due in part to their enhanced expression of HYAL1, whose knockdown restores endothelial stability in PCV. Our key finding explains an intrinsic mechanism of endothelial dysfunction, potentially contributing to leaky choroidal vessels and structurally abnormal vascular dilatation in PCV.

ECM-modifying factors form the central network in our PCV endothelial autocrine mechanism that may drive the hyperpermeability and vessel dilatations observed in PCV eyes. Our comparative single-cell analysis identified increased expressions of ECM-modifiers with established roles in angiogenesis and vascular permeability, which corroborated earlier studies that implicated ECM degradation in PCV pathogenesis [54]. Intriguingly, the heterogeneous flow response in PCV BOECs and differential levels of glycocalyx-related genes led us to hypothesize potential perturbations in the flow-sensing extracellular components of PCV endothelial cells. Glycocalyx, a carbohydrate-rich layer on the luminal surface of vascular endothelium, creates a cell-free, permeable zone between the blood flow and endothelial cells, regulating permeability of the endothelium through size and steric hindrance [55, 56], signaling by plasma-borne endocrine factors and reducing attachment of inflammatory immune cells [57, 58]. The glycocalyx plays an important role in the mechanotransduction of shear stresses. Enzymatic degradation or shear-induced shedding of any component of the glycocalyx can severely impact some of these functions [58, 59]. Infusion of canine femoral arteries with hyaluronidase or cultured endothelial cells with heparitinase both resulted in reduction of shear-induced nitric oxide (NO) production [60, 61]. The application of shear stress to human umbilical vein endothelial cells (HUVEC) also resulted in an increase in HA in the glycocalyx in a postulated positive feedback mechanism [62]. Hyaluronidase-1 is an endocytosed, acid-active enzyme that can break down HA chains of any size into tetrasaccharides [63, 64]. While plasma-borne HYAL1 has been found to have a short half-life of around 2-3 mins [65, 66], we found very low levels of HYAL1 in our PCV vitreous humor samples in contrast to plasma, therefore indicating a likely autocrine role for the increased HYAL1 produced by PCV endothelial cells.

Elevated levels of HYAL1 and HA have also been reported in systemic diseases such as severe dengue [67] and diabetes [68] where glycocalyx degradation, vascular instability and hyperpermeability effects have been associated. Of note, age-related reduction in HA in the Bruch’s membrane have been observed in human eyes [69]. Consistent with reports where short-chain HA deposits can direct and drive endothelial cell migration [68, 70], our data shows that HYAL1 levels can mediate endothelial cell migration. Both increased barrier permeability and endothelial cell migration are processes linked to overall vascular instability. We are mindful that we do not have typical neovascular AMD as a comparison in the current study. Our patient cell-based studies may paint an incomplete picture for the development of aneurysmal dilatations in PCV. Nonetheless, we were able to uncover and demonstrate a previously unreported role for glycocalyx integrity in PCV pathogenesis and present HYAL1 as an autocrine mediator of endothelial dysfunctions in PCV endothelial cells (Fig. 7).

**Figure 7:**
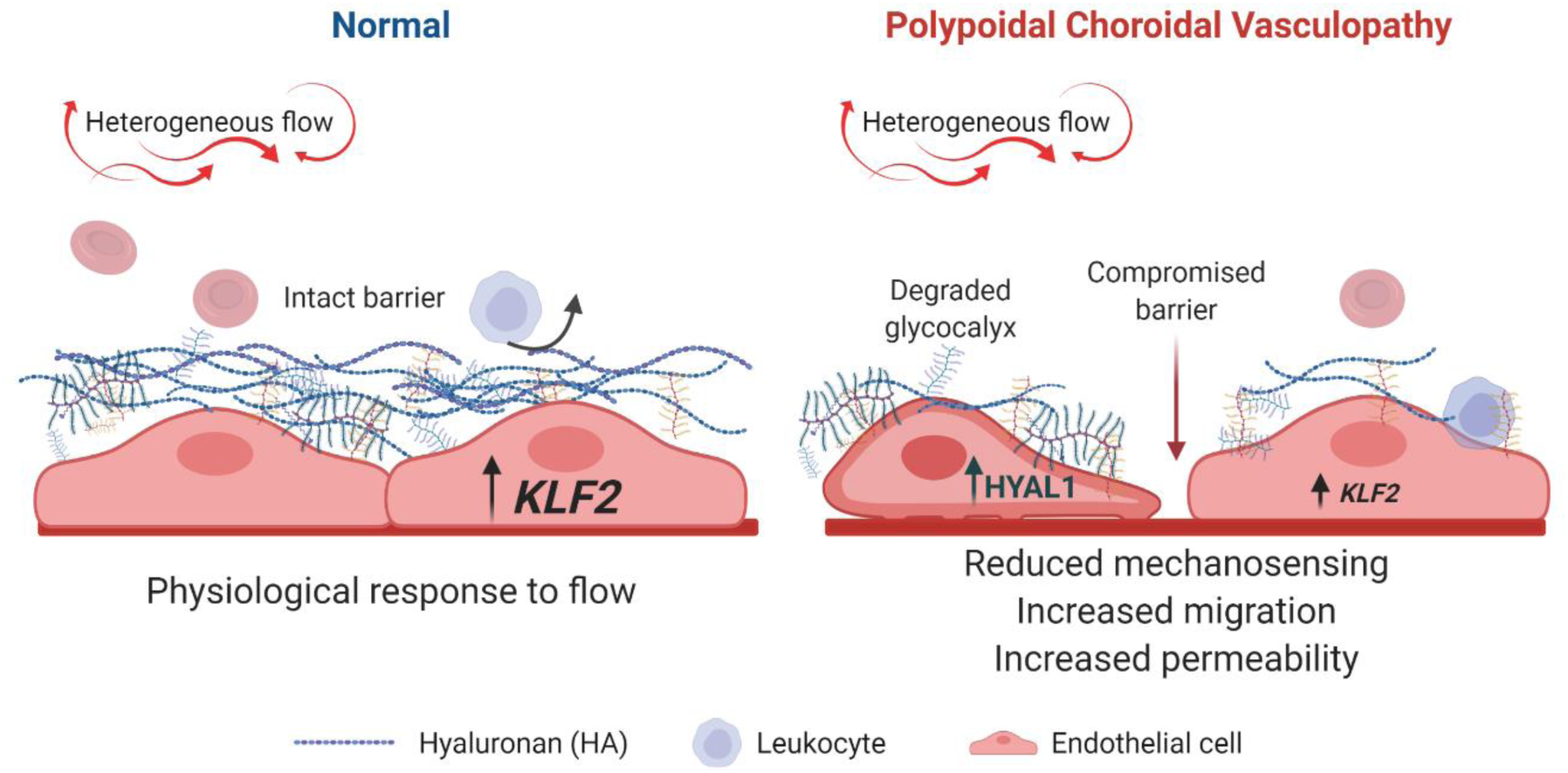
Increased expression of hyaluronidase-1 drives endothelial abnormalities in PCV. Phenotypes indicative of vascular instability were observed in PCV patient-derived endothelial cells under heterogeneous flow conditions. Increased HYAL1 expression in PCV cells led to impaired glycocalyx through the degradation of HA components, giving rise to an altered response to flow, increased cell migration and barrier permeability.

## Conclusions

Understanding mechanisms of diseases affecting ocular integrity is an important area. Our patient endothelial model provides molecular and phenotypic insights into PCV pathophysiological processes which would inform further development of *in vivo* models, as well as pave the way for therapeutic advancement. The fundamental endothelial mechanism presented here could be far-reaching beyond PCV, and potentially a contributor to the pathogenesis of ocular diseases with a vascular origin, suggesting that many pathologies could be ameliorated by better knowledge of endothelial disease biology.

## Materials and Methods

### Patient selection and sample collection

We enrolled subjects from the retina clinic of the Singapore National Eye Center. Inclusion criteria were age 40-80 years (demographics detailed in Supplemental Tables S1 and S2). Written informed consent was obtained from each participant. This study was approved by the Local Ethics Committee of SingHealth Centralised Institutional Review Board (CIRB Refs: R1496 and 2018/2004) and Nanyang Technological University Singapore Institutional Review Board (IRB-2018-01-026 and IRB-2019-03-011-01).

For patients with PCV, the clinical diagnosis was confirmed on fundus examination and fluorescein and indocyanine green angiography. PCV was confirmed based on the presence of polypoidal dilatations on ICGA. Healthy volunteers were recruited for the control group. Controls were further selected based on absence of AMD or PCV from clinical examination. For sample collection, 10mL of fresh blood was collected from each participant and processed in the laboratory within 6 hours. Upon ficoll centrifugation of the blood specimen, a buffy coat layer containing peripheral blood mononuclear cells (PBMCs) was isolated from which DNA extraction was performed for genotyping with the OmniExpress chip. The rest of the PBMCs was used for two purposes - (1) Cultivated in cell culture to derive blood outgrowth endothelial cells (BOECs); (2) Expanded in cell culture and used for reprogramming towards induced pluripotent stem cells (iPSCs).

### Derivation of BOECs and culture conditions

Blood samples were collected from donors as detailed in Supplemental Table S1. BOECs were generated as per described [32] with modifications. Briefly, peripheral blood samples (5 – 9 mL per donor) was diluted 1:1 with phosphate-buffered saline (PBS) and separated to obtain buffy coat by density gradient centrifugation over Ficoll® Paque (GE Healthcare). The buffy coat, which was enriched with peripheral blood mononuclear cells (PBMCs), was carefully collected, washed with PBS, resuspended in heparin-free, EGM-2 medium (Lonza) supplemented with 16% defined foetal bovine serum (FBS; Hyclone) and counted. Plasma was also collected and stored at -80°C. Then, the PBMCs were seeded into collagen I-coated well(s) accordingly so that the cell density was ≥1.5 × 10^6^ cells/cm^2^. Medium was changed every two to three days. Outgrowth colonies should appear between seven to 14 days post-seeding. The cells were expanded to passage 3 before any applications were performed on them, including phenotyping and functional evaluation, in order to opt out unwanted leukocytes. After passage 3, BOECs were cultured on collagen-I-coated tissue culture dishes in heparin-free, EGM-2 with 10% heat-inactivated FBS with media change every 2 – 3 days. BOECs from passages 4 to 8 were used in experiments.

### Endothelial tube formation

Tube formation assay was performed according to manufacturer instructions (Endothelial Cell Tube Formation Assay, Corning). More details are found in Supplemental Methods.

### Fibrin gel bead sprouting assay

To evaluate the angiogenesis ability of BOECs, fibrin gel bead sprouting assay was performed as per described [71] with modifications. More details are found in Supplemental Methods.

### Induced pluripotent stem cell generation and RPE cell differentiation

Frozen PBMCs that were previously obtained from density gradient centrifugation via Ficoll-Paque, were thawed and reprogrammed using CytoTune-iPS 2.0 Sendai Reprogramming Kit (Thermo Fisher Scientific) under Reprogram PBMCs (Feeder-free) section. Once emerging colonies were observed, they were manually picked for expansion into individuals iPS cell lines while being transferred to Matrigel-coated plates (Corning, 354234) and cultured in mTeSR™1 (STEMCELL Technologies). After successfully stabilizing of at least 2 clones per reprogrammed cell line, cells were expanded further in mTeSR™1 and passaged using 0.5mM EDTA passaging solution (Sigma Aldrich).

Patient-specific iPS cell lines undergo RPE differentiation following protocol [34] with some modifications. Changing of medium with the addition of growth factors was done on day 0, 3,5 and 7 then every 2 days till first pigmentation was observed (day 20). Around Day 30, immature RPE cells were seeded at a density of 1 × 10^5^ cells/cm^2^ onto growth factor reduced ECMH (Corning)-coated transwells and allowed to be mature for 4-6 weeks in RPE medium (XVIVO 10, Lonza). Matured RPE at P1 or P2 were used in downstream experiments.

### Functional characterization of iPSC-RPE cells

More details are found in Supplemental Methods.

### Orbital flow setup

BOECs were trypsinized and counted with 0.4% Trypan Blue (Gibco, Thermo Fisher Scientific) staining on an automated cell counter (Countess II, Thermo Fisher Scientific) before diluting and seeding onto rat-tail collagen Type1 (Corning)-coated 12-well plates to give 120,000 viable cells per well. Seeded wells were incubated for 48h in a 5% CO_2_, 37°C, humidified incubator before overlying media was replaced with 800μl of fresh heparin-free, EGM-2 medium (10% heat-inactivated FBS) to give a liquid height of ∼2mm per well. Plates were then replaced onto an orbital shaker in a 5% CO_2_, 37°C, humidified incubator and rotated at 210rpm for 24h. Static controls were setup in the same manner under the same conditions, with 800μl of fresh heparin-free, EGM-2 medium (10% heat-inactivated FBS) replaced after 48h post-seeding and replaced into the incubator without rotation.

### Single-cell RNA sequencing and analysis

BOECs from 2 PCV and 2 normal lines were seeded and subjected to orbital flow as described in Orbital Flow Setup. After 24h of orbital flow, the BOECs were trypsinized, resuspended in heparin-free, EGM-2 medium (10% heat-inactivated FBS) and counted using Trypan blue and an automated cell counter (Countess II, Thermo Fisher Scientific) and resuspended appropriately for loading onto 10X Genomics Chromium Controller chip by facility personnel at Single-cell Omics Centre (SCOC), Genome Institute Singapore (GIS). Each BOEC cell line was prepared as a separate scRNA-seq library using Chromium Single Cell 3’ v3 Reagent Kit (10X Genomics) by SCOC GIS and the final ready-to-sequence libraries were handed over with quantification and quality assessment reports from Bioanalyzer Agilent 2100 using the High Sensitivity DNA chip (Agilent Genomics). Individual libraries were pooled equimolarly and sent for sequencing by NovogeneAIT Genomics (Singapore). Raw sequencing data was also processed by NovogeneAIT Genomics (Singapore) using CellRanger (10x Genomics) with reads mapped to the human genome assembly (GRCh38).

We performed secondary analysis on the resultant filtered matrix files using Seurat (v 3.2.0) [72]. Data was filtered for dead/poor quality cells based on low number of genes detected (<200) or potential doublets (>9500) as recommended by Seurat’s tutorial (satijalab.com) and inspection of nFeature spread for each sample (Supplemental Fig. S2a). Cells with high percentage of mitochondrial genes were also removed with the threshold of less than 20% informed by a previously reported percentage of mitochondrial gene content in endothelial cells [73]. In order to inspect for any cell cycle heterogeneity between samples, cell cycle states for each sample were determined by the *CellCycleScoring* function in Seurat (Supplemental Fig. S2b). These filtered datasets were then scaled and normalized using *SCTransform* individually before integrated based on 3000 integration features. Clusters were identified in the integrated dataset using the *FindCluster* function at resolution 0.2, after PCA analysis, *RunUMAP* and *FindNeighbours* at 1:30 dimensions. Marker genes were then identified for each cluster using *FindAllMarkers* with MAST [74] (R package) as the selected test of choice. Different expression analysis between PCV and normal datasets were performed using *FindMarkers* with MAST for each individual cluster. Gene enrichment analysis was carried out for both marker genes of clusters and differential expression genes between PCV and normal in each clusters using clusterProfiler (R package, v 3.17.0.) [75]. *AddModuleScore* function (Seurat) was used to present overall relative expression profiles between PCV and normal, for genes found in the indicated enriched processes.

### MitoSOX assay

BOECs treated to 24h of flow or static conditions on glass-bottom wells were stained with MitoSOX Red mitochondrial superoxide indicator according to manufacturer instructions (Cat. no. M36008, Thermo Fisher Scientific). More details are found in Supplemental Methods.

### Mitochondrial function assay by Seahorse analyzer

BOECs were reseeded onto collagen-I-coated 96-well Seahorse microplates at 20,000 cells per well and incubated in a humidified incubator at 37°C, 5% CO_2_ for 5h. Thereafter, they were prepared and assayed for mitochondrial function assessment according to manufacturer instructions (Seahorse XF Cell Mito Stress Test Kit, Agilent Technologies). More details are found in Supplemental Methods.

### Wound healing assay

BOEC cultures, prepared as described in Orbital Flow Setup section, were scratched across the horizontal diameter of each well with a P200 micropipette tip before imaging on an automated microscope (Celldiscoverer 7, ZEISS) using a 5x objective over time. Multi-tile images were captured to encompass the entire scratch wound at time=0h, under live-cell imaging conditions (5% CO_2_, 37°C). Culture plates were then replaced into a 5% CO_2_, 37°C, humidified incubator and left to recover without rotation (for both rotated and static conditions) for 21h. The same tiling positions determined for each well at 0h were reused for the imaging at 21h. Wound area was obtained using manual region-of-interest (R.O.I) annotation of cell free areas across each tiled image set before area of R.O.I. was obtained using Measure in Fiji [76]. Wound closure percentage shown is calculated by taking the difference between cell-free areas for t=0h and t=16h and dividing it by t=0h % per well. All data points were collected over a total of 3 independent experiments.

### Permeability imaging assay

BOEC cultures were prepared as described in Orbital Flow Setup section with the exception of biotinylated gelatin coating replacing rat-tail collagen coat. Biotinylated gelatin was prepared as described [77] and plates were coated with 10mg/ml of biotinylated gelatin diluted in 0.1M sodium bicarbonate solution (pH8.3) at 4°C overnight. Solutions were removed and wells washed with PBS before preparing cultures as describe in Orbital Flow Setup section. After 24h of rotated/static incubation, culture plates were left to recover without rotation (for both rotated and static conditions) for 21h in a 5% CO_2_, 37°C, humidified incubator. After which overlying medium was removed, rinsed with PBS, before staining with 2μg/ml FITC-Neutravidin (A2662, Thermo Fisher Scientific) for 3mins in a 5% CO_2_, 37°C, humidified incubator. Stained wells were washed thrice with PBS before they were fixed with 4% paraformaldehyde phosphate-buffered solution (Nacalai Tesque) for 10mins at room temperature. Fixed cell layers were washed with PBS once before automated imaging using a 5x objective (CellDiscoverer7) to capture a tiled image of each entire well. FITC stained areas per well were obtained using ZEN BLUE Image Analysis tool’s interactive Segmentation module. Relative FITC area was determined by normalizing total area per well against the average area obtained across all normal BOEC lines used in that experiment. All data points were collected over a total of 3 independent experiments.

### Hyaluronidase activity assay

Hyaluronidase activity from rotated conditioned media was measured using an ELISA-like assay described in Lokeshwar *et al.* (2001) [78] with modifications. As shown in the graphical workflow (Fig. 5b, top panel), high molecular weight (1.5M-1.75M) HA (63357, Sigma Aldrich) was coated onto black, clear-bottom 96-well plates at a concentration of 500μg/ml overnight at 4°C. HA solutions were removed and wells were washed with PBS twice. Each HA-coated well was incubated with 10μl of conditioned media and 90μl of HAase assay buffer (0.1M sodium formate, 0.15M NaCl, 0.2mg/ml BSA, pH4.2) at 37°C for 24h. Incubation solutions were removed and wells washed with PBS twice before incubated with 2μg/ml biotinylated-HABP (PBS, 1% BSA) for 30mins at room temperature. Staining solution was removed and wells washed twice with PBS. PBS was replaced at 100μl per well before readout at 490nm excitation and 525nm emission (fixed gain of 150) on a Synergy H1 plate reader (Biotek). Conditioned media from at least 2 independent experiments per cell line were analyzed and shown.

### Glycocalyx HA staining

Rotated BOEC cultures were prepared as indicated in Orbital Flow Setup with the exception of glass-bottom 12-well plates (Cellvis) replacing polypropylene 12-well plates used in other experiments. Overlying medium was removed and cell monolayers were briefly but gently washed once with cold sterile PBS once before fixation with cold methanol for 10mins at -20°C. Fixed cell layers were washed gently with PBS once before addition of endogenous biotin blocking solutions (Endogenous Biotin Blocking kit, Thermo Fisher Scientific) according to manufacturer’s instructions. Biotin and avidin-blocked cell layers were then washed with PBS and stained with 1μg/ml of biotinylated-HABP (versican G1 domain, Affirmus Biosource) diluted in PBS at 4°C overnight. Cell layers were then washed twice with PBS before counterstaining with Hoechst 33342 (2 drops per ml, Ready Flow, Thermo Fisher Scientific) and 20μg/ml FITC-Neutravidin (A2662, Thermo Fisher Scientific) for 30mins. Staining solution was removed and cells washed with PBS before z-stack imaging using confocal microscopy. Start of z-stacks was determined using Hoechst 33342 signal staining for nuclei and last positions were determined by the last visible FITC signal per sample. 3 frames per region per cell line were imaged using a 40x objective. FITC stained areas per frame were obtained and summed across all stacks using ZEN BLUE Image Analysis tool’s interactive Segmentation module.

### siRNA knockdown

siRNA against human HYAL1 (ON-TARGETplus SMARTpool, Dharmacon) and a non-targeting control (ON-TARGETplus NT#4, Dharmacon) were prepared in 1x siRNA buffer (Dhamarcon) according to manufacturer’s instructions to give a stock concentration of 20μM. Employing reverse transfection, siRNA and transfection reagent (Dharmafect-1, Dharmacon) were first complexed together in serum-free heparin-free, EGM-2 medium for 20mins at room temperature before adding into empty, collagen-coated plates. BOECs were then prepared as described in Orbital Flow Setup and seeded into each well to give a final siRNA concentration of 25 or 50nM and 2.5μl of Dharmafect-1 per well. Experiments proceed as described in Orbital Flow Setup, Wound Healing Assay or Permeability Imaging Assay.

### Confocal and automated microscopy imaging

All confocal imaging were carried out at NTU-Optical Bio-Imaging Centre on an Inverted Confocal Airyscan Microscope (LSM800, ZEISS) and automated imaging was carried out on the CellDiscoverer7 (ZEISS).

## Statistics

Data analysis (excluding all single-cell RNA sequencing analyses) was performed with GraphPad Prism version 9.0.2. Data were tested for normality using the D’Agostino & Pearson test or Shapiro-Wilk test. Where necessary, data that failed normality tests were log-transformed before statistical analysis. *P* values for data with a single factor were obtained using a two-tailed *t*-test (parametric) or two-tailed Mann-Whitney test (non-parametric) as indicated. *P* values for data with 2 factors were assessed using a two-way ANOVA with Tukey’s multiple comparisons test. A value of *P* < 0.05 was considered statistically significant. Other relevant statistical considerations have been elaborated in figure legends.

## List of abbreviations

AMD: age-related macular degeneration
BOEC: blood outgrowth endothelial cell
GWAS: genome-wide association studies
iPSC: induced pluripotent stem cell
PBMC: peripheral blood mononuclear cell
PCV: polypoidal choroidal vasculopathy
RPE: retinal pigmented epithelial
siRNA: small-interfering RNA
VEGF: anti-vascular endothelial growth factor

## Declarations

### Ethics Approval

This study was approved by the Local Ethics Committee of SingHealth Centralised Institutional Review Board (CIRB Refs: R1496 and 2018/2004) and Nanyang Technological University Singapore Institutional Review Board (IRB-2018-01-026 and IRB-2019-03-011-01).

### Consent to participate

Informed consent was obtained from all individual participants included in the study.

### Consent for publication

Informed consent was obtained from all individual participants. There is no personally identifiable information in this article.

## Acknowledgements

We thank all patients and healthy donors who have participated in this study. Special thanks to Ms Kelly Wong Kai Li and Dr Srivani Sistla for coordinating clinical sample collection, Dr Shyam Prabhakar and Dr Samydurai Sudhagar for advice on 10x Genomics single cell library preparations, Dr Balakrishnan Kannan for his guidance in confocal and widefield microscopy imaging, A/Prof Yusuf Ali and his team for the lending of laboratory space to run Seahorse Analyzer experiments, Dr Anthony Siau for helping with experimental optimization, and LKCMedicine Research Administration and Support Services for their assistance.

## Author contributions

Conceptualization: CC

Data curation: KXW, NJYY, CYN, FWJC, FQ

Formal analysis: KXW, NJYY, CC

Investigation: KXW, NJYY, CYN, FWJC, FQ

Methodology: KXW, NJYY, CYN, FWJC, FQ, XFT, YBX, GN, HMT

Resources: CC, CMGC, HWH

Validation: KXW, NJYY

Visualization: KXW, NJYY, CYN, CC

Funding acquisition: CC, CMGC, XS

Project administration: CC, CMGC

Supervision: CC, CMGC, XS, NRD, HWH

Writing – original draft: KXW, CC

Writing – review & editing: All authors

All authors approved the manuscript.

## Availability of Data and Materials

Further information and requests for resources and reagents should be directed to and will be fulfilled by the lead contact, Christine Cheung. Most materials used in this study are commercially procured. There are restrictions to the availability of blood outgrowth endothelial cell lines derived from human patients and normal donors due to ethics considerations for use of these materials within the current scope of study. Requests can be made to the lead contact as we will explore use of materials subject to new ethics approval and research collaboration agreement (including material transfer).

The authors declare that all data supporting the findings of this study are available within the paper and supplemental information that includes original data of western blots. Specifically, single-cell sequencing (scRNA-seq) dataset that support the findings of this study are available upon request from the corresponding author or will be available in public repository when this manuscript is published at refereed journal.

---

**Supplemental Figure S1:**
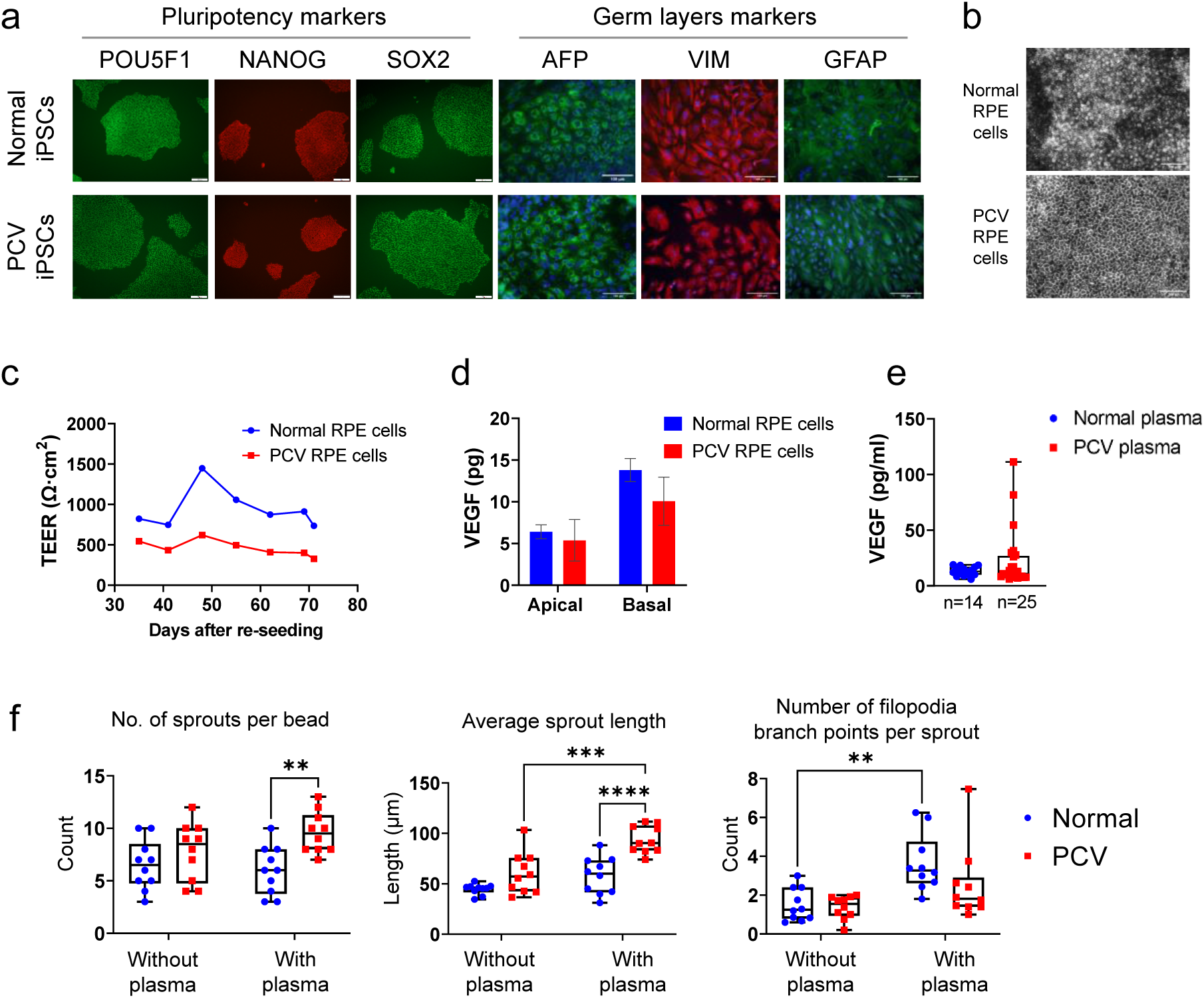
Extrinsic mediators influence sprouting angiogenesis of BOECs. **(a)** Representative images of immunostaining for pluripotency and germ layer markers on PCV and normal donor-derived iPSCs. Scale bar, 100 um. **(b)** RPE cells differentiated from PCV and normal donors developed pigment from day 34. Scale bar, 100 um. **(c)** Transepithelial electrical resistance (TEER) assay on iPSC-RPE cells between 34-71 days post-seeding onto transwells to establish RPE polarity. Ave Ω of each RPE cell line was calculated from 3 independent transwells, 3 TEER readings/ transwell. **(d)** ELISA measurement of VEGF in conditioned media collected from the apical and basal compartments in RPE cell transwell cultures. Conditioned media of 4 PCV and 4 normal iPSC-RPE cell lines were analysed at day 86 of differentiation. **(e)** ELISA measurement of VEGF from plasma samples of 25 PCV and 14 normal donors. Statistical tests were assessed using two-tailed t-tests with Welch’s correction but no significant differences between Normal and PCV were observed. **(f)** BOECs were coated onto microcarrier beads and allowed to sprout either in standard culture media (EGM-2 with 10% heat-inactivated FBS) or media containing isogenic patient plasma (EGM-2 with 10% plasma) for 24 hours in a fibrin sprouting assay. Quantification of number of sprouts per bead, average sprout length and number of filopodia branch points per sprout was performed using Imaris; n = 10 beads per donor cell line. Data are presented as box and whiskers plots showing the median, minimum, maximum and quartiles of each group. *P* values between individual groups were obtained using two-way ANOVA with Tukey’s multiple comparisons test. \*\**P*<0.01, \*\*\**P*>0.001.

**Supplemental Figure S2:**
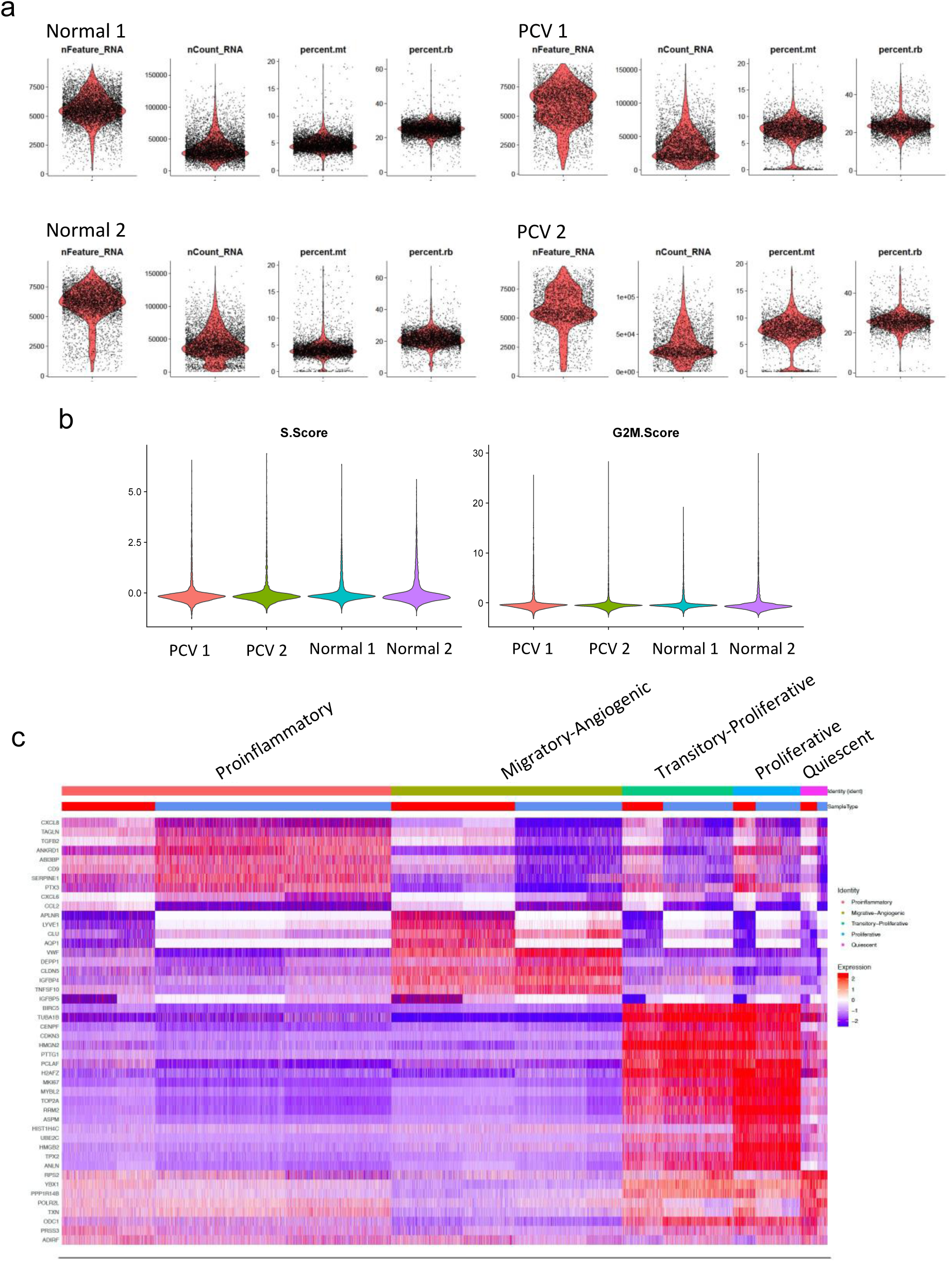
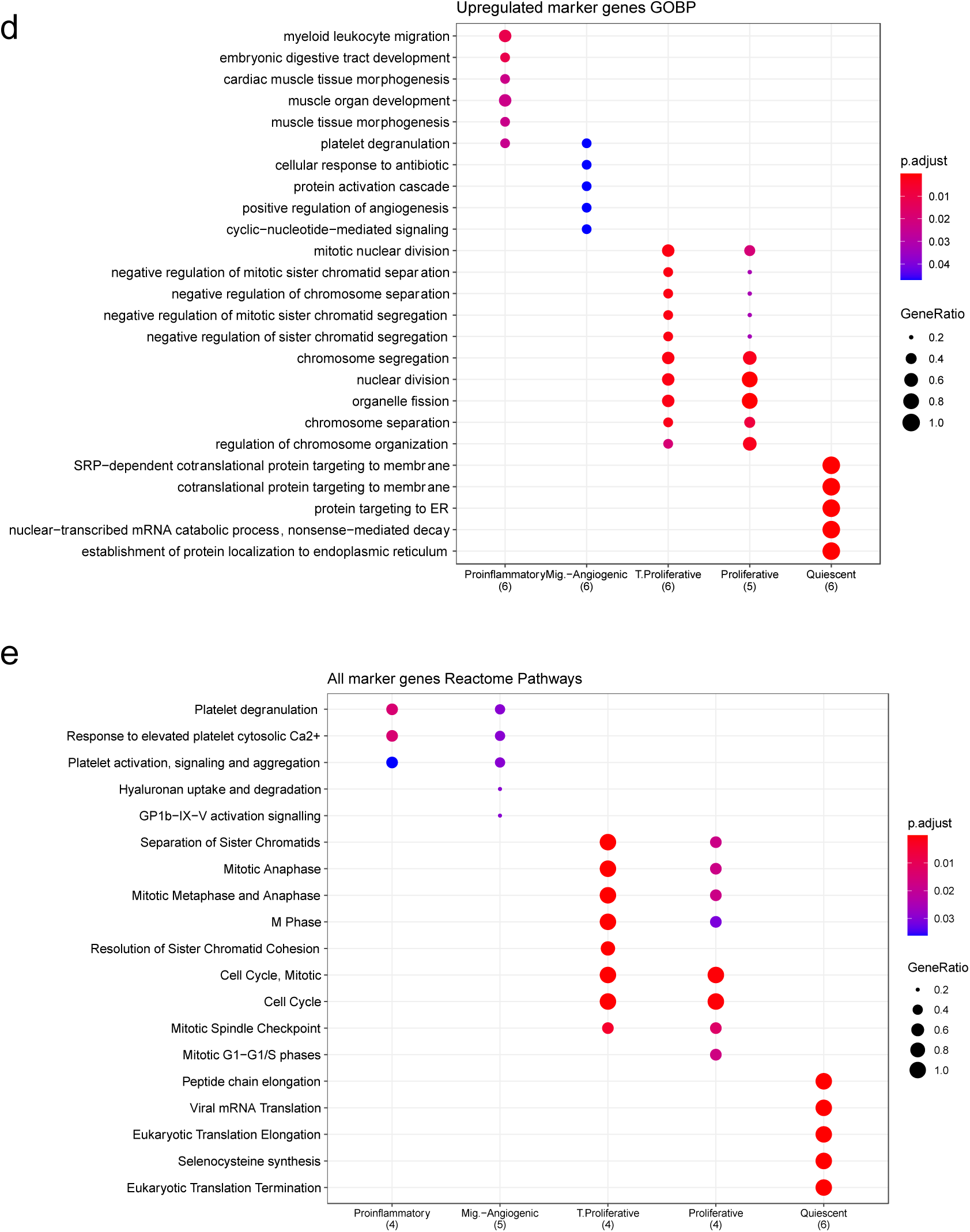
scRNA-seq data quality checks and cluster marker enrichments. **(a)**.Violin plots with dots representing individual cells for the number of features, number of counts, percentage of mitochondrial genes (mt) and percentage of ribosomal genes (rb) found in each individual library **(b)** Violin plots showing expression scores for S-phase and G2M genes in each individual library. **(c)** Heatmap showing expression of top 10 marker genes for each clusters at the single cell level. Dotplots of the top unique gene sets, GOBP in **(d)** and Reactome in **(e),** enriched for the upregulated marker genes of each identified cluster shown here ranked by Bonferroni corrected *p*-values. Right panel shows violin plots that represent calculated module scores for the processes indicated.

**Supplemental Figure S3:**
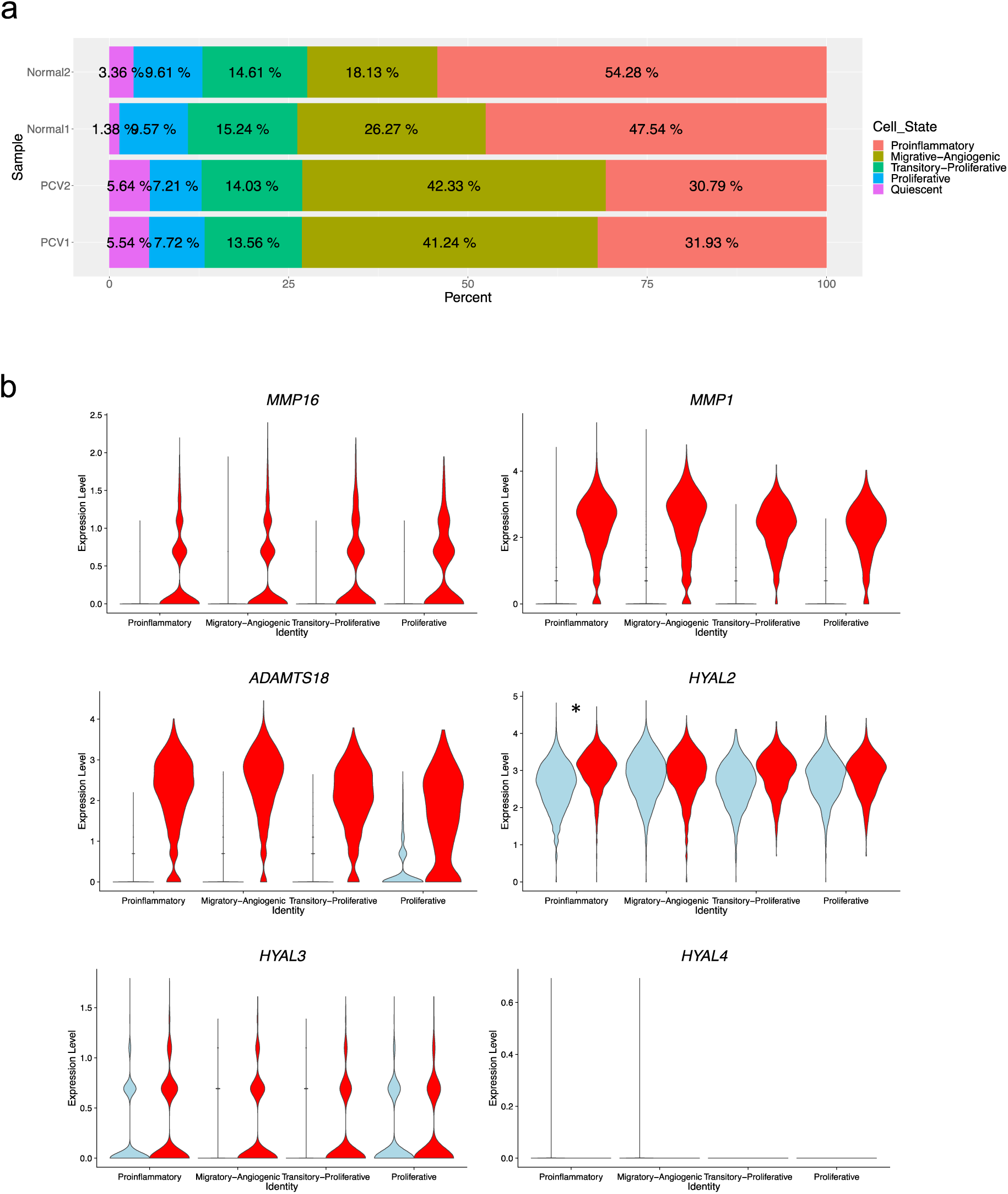
Differential expression in PCV and normal BOECs after orbital flow. **(a)**. Percentage breakdown of cells per cell state for individual PCV and normal libraries. **(b)** Violin plots showing differential expression of ECM-modifying genes and other hyaluronidases detected in the study, *adj. *p*< 0.01.

**Supplemental Table S1:** Demographics details of PCV patients and normal individuals.

| Disease status | Age | Blood outgrowth endothelial cells (BOECs) | hiPSC-derived retinal pigment epithelial cells (iPSC-RPE cells) | Genotypes |  |  |
| --- | --- | --- | --- | --- | --- | --- |
|  |  |  |  | ARMS2 rs10490924 (risk allele: T) | HTRA1 rs11200638 (risk allele: A) | CFH rs800292 (risk allele: G) |
| PCV | 71 | ✓ | ✓ (2 iPSC clones) | TT | AA | GA |
|  | 86 | ✓ |  | TT | AA | AA |
|  | 73 |  | ✓ (2 iPSC clones) | TT | AA | GA |
|  | 71 | ✓ |  | TT | AA | GG |
|  | 71 | ✓ |  | GT | GA | GA |
| Control | 46 | ✓ | ✓ (2 iPSC clones) | GT | GA | GG |
|  | 46 | ✓ | ✓ (2 iPSC clones) | GG | GG | GA |
|  | 71 | ✓ |  | GG | GG | GG |
|  | 49 | ✓ |  | GT | GA | GA |
|  | 47 | ✓ |  | GT | GA | GG |
|  | 46 | ✓ |  | GT | GA | GA |

**Supplemental Table S2:** HYAL1 validation in clinical samples.

| Characteristics, N (%) | Plasma of PCV patients (n = 25) | Vitreous humor of PCV patients (n = 17) |
| --- | --- | --- |
| Age* | 66.8 (11.5) | 69.9 (8.1) |
| Age, Male* | 67.3 (11.9) | 71.2 (9) |
| Age, Female* | 65.2 (11.3) | 67.7 (6.3) |
| Gender, male | 19 (76) | 6 (35.3) |
| Gender, female | 6 (24) | 11 (64.7) |
| Ethnicity | Chinese: 25 (100) | Chinese: 13 (76.4)<br>Malay: 2 (11.8)<br>Indian: 1 (5.9)<br>Filipino: 1 (5.9) |

All values are reported as N (%) where N indicated number of observations.
\*Values are expressed as mean (± standard deviation).

**Supplemental Table S3:** Key resources used in this study.

| REAGENT or RESOURCE | SOURCE | IDENTIFIER |
| --- | --- | --- |
| **Antibodies** |  |  |
| Mouse monoclonal anti-Oct-3/4 | Santa Cruz | Cat# sc-5279,<br>RRID:AB_628051 |
| Mouse monoclonal anti-Sox2 | R&D systems | Cat# MAB2018,<br>RRID:AB_358009 |
| Goat polyclonal anti-Nanog | R&D systems | Cat# AF1997,<br>RRID:AB_355097 |
| Mouse monoclonal anti-AFP | Sigma-Aldrich | Cat# A8452,<br>RRID:AB_258392 |
| Mouse monoclonal anti-β-III-Tubulin | Covance | MMS-435P,<br>RRID:AB_2313773 |
| Chicken polyclonal anti-Vimentin | Merck Millipore | Cat# AB5733,<br>RRID:AB_11212377 |
| Mouse monoclonal anti-CD31-APC (clone wm59) | BioLegend | Cat# 303116,<br>RRID:AB_1877151 |
| Mouse monoclonal anti-CD144-PE (clone 55-7H1) | BD Pharmingen | Cat# 560410,<br>RRID:AB_1645502 |
| Mouse monoclonal anti-CD45-APC (clone 2D1) | Invitrogen | Cat# 17-9459-42,<br>RRID:AB_10718532 |
| Mouse monoclonal anti-CD68-FITC (clone Y1/82<sup>a</sup>) | BioLegend | Cat# 333806,<br>RRID:AB_1089054 |
| Mouse monoclonal anti-CD133-APC (clone 7) | BioLegend | Cat# 372806,<br>RRID:AB_2632882 |
| Rabbit polyclonal anti-HYAL1 | Thermo Fisher Scientific | PA5-79420,<br>RRID:AB_2746536 |
| Rabbit polyclonal anti-beta-Actin | Abcam | Cat# ab75186,<br>RRID:AB_1280759 |
| Mouse monoclonal anti-CAV1 | Thermo Fisher Scientific | Cat# MA3-600,<br>RRID:AB_779568 |
| Goat polyclonal anti-CDH5 | Santa Cruz Biotechnology | Cat# sc-6458,<br>RRID:AB_2077955 |
| **Biological samples** |  |  |
| PBMC | This paper | N/A |
| Plasma | This paper | N/A |
| Vitreous humor | This paper | N/A |
| **Chemicals, peptides, and recombinant proteins** |  |  |
| Biotinylated-HABP (versican G1 domain) | Amsbio | AMS.HKD-BC41 |
| TRITC-Phalloidin | Merck | 90228 |
| MitoSOX Red | Thermo Fisher Scientific | M36008 |
| **Critical commercial assays** |  |  |
| CytoTune-iPS 2.0 Sendai Reprogramming Kit | Thermo Fisher Scientific | A16517 |
| Chromium Single Cell 3' v3 Reagent Kit | 10X Genomics | PN-1000075 |
| Seahorse XF Cell Mito Stress Test Kit | Agilent Technologies | 103015-100 |
| VEGF ELISA | Abcam | ab100662 |
| PEDF ELISA | Abcam | ab213815 |
| HTRA1 ELISA | CUSABIO | CSB-EL010901HU |
| HYAL1 DuoSet ELISA and DuoSet ELISA Ancillary Reagent Kit 2 | R&D Systems | DY7358 and DY008 |
| **Experimental models: Cell lines** |  |  |
| Patient-derived BOECs | This paper | N/A |
| Patient-derived iPSCs | This paper | N/A |
| iPSC differentiated RPE | This paper | N/A |
| Oligonucleotides |  |  |
| Non-targeting siRNA (ON-TARGETplus NT#4) | Dharmacon | D-001810-04-05 |
| siRNA pool for HYAL1 (ON-TARGETplus SMARTpool) | Dharmacon | L-010516-00-0005 |
| Software and algorithms |  |  |
| Seurat v 3.2.0 | 1 | https://satijalab.org/seurat/ |
| clusterProfiler v 3.17.0 | 2 | https://github.com/YuLab-SMU/clusterProfiler |
| ImageJ/Fiji | 3 | https://imagej.net/Fiji/Downloads |
| ZEN BLUE | ZEISS | https://www.zeiss.com/microscopy/int/products/microscope-software/zen.html |
| FlowJo | Becton Dickinson | https://www.flowjo.com/ |
| Quest Graph™ Four Parameter Logistic (4PL) Curve Calculator | AAT Bioquest, Inc | https://www.aatbio.com/tools/four-parameter-logistic-4pl-curve-regression-online-calculator |
| Prism version 9.0.2. | GraphPad | https://www.graphpad.com/scientific-software/prism/ |
| Imaris 3.0 | Oxford Instruments | https://imaris.oxinst.com/ |
| Other |  |  |
| EVOM2 epithelial voltohmmeter | World Precision Instruments | N/A |
| Synergy H1 | BioTek | N/A |
| Cytation 3 | BioTek | N/A |
| Confocal Airyscan Microscope LSM800 | ZEISS | N/A |
| CellDiscoverer7 | ZEISS | N/A |
| BD LSR Fortessa X-20 cell analyser | Becton Dickinson | N/A |

## Supplemental Methods

### Endothelial tube formation

Tube formation assay was performed according to manufacturer instructions (Endothelial Cell Tube Formation Assay, Corning). BOECs were seeded onto Matrigel as 20,000 cells per well of a 96-well plate in serum-free EGM-2 medium, and H9 embryonic stem cell line (H9-ESC) and human umbilical vein endothelial cells (HUVEC) were seeded 7,500 cells per well in mTeSR medium (STEMCELL Technologies) and serum-free, heparin-free, EGM-2 medium, respectively. Phase-contrast micrographs (1024 × 1024-pixel) of tubular networks were captured at 2, 4, 6 and 8h using a Nikon Ti-E inverted microscope at 4× magnification and image acquisition was performed using MetaMorph version 7.8 (Molecular Devices). ImageJ was used to crop four different 380 × 380-pixel areas (optical fields) from each original 4× image. Thereafter, tube formation parameters were quantified using Angiogenesis Analyzer on ImageJ.

### Fibrin gel bead sprouting assay

To evaluate the angiogenesis ability of BOECs, fibrin gel bead sprouting assay was performed as per described ^4^ with modifications. Briefly, BOECs were coated onto Cytodex 3 microcarrier beads (Sigma-Aldrich) at 150 cells/bead with agitation for 4h and allowed to adhere overnight. Coated beads were subsequently suspended in fibrinogen solution at a concentration of 500 beads/mL and clotted with thrombin. Gels were then topped up with heparin-free, EGM-2 with 10% FBS (Gibco). Cells were incubated overnight and observed for sprout formation after 24h.

Cells were incubated for 24h and gels containing sprouts were fixed with 4% paraformaldehyde (200 µL/well) overnight at 4°C. Gels in 8-well chamber slides were washed by rinsing twice in 100 µL/well 1×PBS and gentle orbital shaking (agitation) in 1×PBS for 30 min. Permeabilization was performed with agitation in 0.5% Triton X-100 for 20 min and staining with agitation in TRITC-Phalloidin in 1×PBS with 1% BSA for 30 min and 500 ng/ml DAPI for 10 min. Final rinses were performed with agitation in PBS/T (0.1% v/v Tween-20 in 1×PBS) and subsequently in 1×PBS. Stained gels were stored in 1×PBS at 4°C before confocal imaging.

Fibrin-embedded BOECs were imaged using an inverted laser scanning confocal microscope (LSM800, Carl Zeiss) using a Plan-Apochromat 20x/0.80 objective lens. Two-channel Z-stack images (AF568 and DAPI) of whole beads were captured using the ZEN software (blue edition, Carl Zeiss). Images of 1024 × 1024-pixel resolution were acquired from 0.6× optical zoom at Z-intervals of 1.11 µm. Approximately 100 – 200 Z-slices were acquired for each bead. For each individual, 2 – 4 of the most well-formed individual filopodia were imaged. Sprouting parameters were quantified using Imaris 3.0 (Oxford Instruments). The number of sprouts per bead, ‘Filament Tracer’ sprout length and number of filopodia branch points per sprout were measured.

### Endothelial phenotyping by flow cytometry

Cell surface markers were quantified using flow cytometry to phenotypically confirm the endothelial identity of the derived BOECs. CD31 and CD144 were selected as endothelial markers; CD45 and CD68 as leukocyte exclusion markers, and CD133 as a progenitor marker. Briefly, the BOEC monolayers were trypsinised and washed with DPBS prior to staining with the marker antibodies in the dark for 15 min, room temperature. Fluorescence data were collected on a BD LSR Fortessa X-20 cell analyser (Becton Dickinson) and analyzed using FlowJo software (Becton Dickinson). Gating strategy for FSC/SSC plots and positive/ negative staining have been presented in Supplemental Information.

### Immunostaining

Cells were fixed with 4% Paraformaldehyde Phosphate (09154-85, Nacalai Tesque) at room temperature for 20 minutes, then washed with DPBS without Ca and Mg (SH3002803, Hyclone) and stored at 4°C. Before staining, cells were permeabilized with 0.2% Triton X-100 (Sigma) and blocked with blocking buffer (4% FBS in DPBS) for 60 minutes. Cells were then incubated overnight at 4°C with primary antibodies (Supplemental Table S3) diluted in blocking buffer. Next day, cells were washed three times with wash buffer (TBS + 0.05% Tween20 in water) before incubating with secondary antibodies diluted in blocking buffer for 1 hour at room temperature. Cells were then washed three times with wash buffer followed by keeping in Hoechst dye (1:10000 in PBS). Immunocytochemistry was analyzed using an Olympus IX71 inverted fluorescence microscope fitted with an Olympus digital camera.

### Functional characterization of iPSC-RPE cells

RPE cells were seeded at a density of 1.07×10^5^ cells/cm^2^ in 24well or 6well Transwells (0.4μm pores; Corning) which are coated with 1X Matrigel (Corning). Media had been changed twice per week until cells were confluent. Then an EVOM2 (World Precision Instruments; Sarasota, FL, USA) epithelial voltohmmeter, was used to detect cell–cell resistance every 7 days for continuous 8 weeks. TEER values were subtracted by background resistance and corrected with the culture-insert differences.

iPS-differentiated RPE cells were seeded at a density of 5×10^5^ cells/well in 24mm diameter polyester inserts (0.4μm pores; 3450, Corning) which are coated with 1X Matrigel (Corning). Cells had been cultured for 2-3 months with media changes (XVIVO 10, Lonza) twice per week. Conditioned Medium was collected from both apical and basal chambers 24 hours after feeding cells. ELISA for vascular endothelial growth factor (VEGF; ab100662, abcam), pigment epithelium-derived factor (PEDF; ab213815, abcam), and HtrA Serine Peptidase 1 (HTRA1, CSB-EL010901HU, CUSABIO) were conducted according to the manufacturer’s protocols. Growth factor concentrations were calculated from standard curves and corrected with chamber volumes.

### MitoSOX assay

BOECs treated to 24h of flow or static conditions on glass-bottom wells were stained with MitoSOX Red mitochondrial superoxide indicator according to manufacturer instructions (Cat. no. M36008, Thermo Fisher Scientific). MitoSOX Red-stained cells were fixed with 4% PFA for 15 min and nuclei were stained with DAPI for 10 min. Cells were imaged using an confocal microscopy with a Plan-Apochromat 20x/0.80 objective lens. Z-stacks of 2987 × 2987-pixel resolution were acquired from 1.0× optical zoom at Z-intervals of 1.25 µm. Cells were imaged at the center and the periphery of wells with 3 – 5 representative images captured for each region. Prior to image analysis, image stacks were split into single-channel images and Z-projected with maximum intensity projection using ImageJ. Image analysis was then performed using CellProfiler (version 4.0, Broad Institute) ^5^ with an optimized pipeline to identify whole cells using both nuclei and MitoSOX Red staining followed by measurements of integrated intensity per cell.

### Mitochondrial function assay by Seahorse analyzer

BOECs were reseeded onto collagen-I-coated 96-well Seahorse microplates at 20,000 cells per well and incubated in a humidified incubator at 37°C, 5% CO_2_ for 5h. Thereafter, they were prepared and assayed for mitochondrial function assessment according to manufacturer instructions (Seahorse XF Cell Mito Stress Test Kit, Agilent Technologies). The drugs oligomycin, carbonyl cyanide-4 (trifluoromethoxy) phenylhydrazone (FCCP) and rotenone/ antimycin A (Rot/AA) were used in the assay at concentrations of 1 µM, 1.5 µM and 0.5 µM respectively. To obtain post-assay cell counts for normalization, assayed cells were rinsed once with 1×DPBS, fixed with 4% PFA, stained with DAPI (500ng/ml) for 10 min and counted automatedly using an imaging reader with the Gen5 software (Cytation 3, BioTek Instruments). Thereafter, OCR values were normalized to these counts in individual wells, processed and analyzed in Wave 2.6.1 software according to manufacturer instructions (Agilent Technologies). Wells containing uneven cell distribution or displaying outlier OCR were excluded from analysis.

### Western Blot

Cell lysates were prepared by lysing PBS-rinsed cell layers with 150μl of 1X LDS buffer (NuPAGE, Thermo Fisher Scientific) at 4°C for 10mins with gentle rocking. Lysates were harvested and heated at 95°C for 10mins before being resolved in 10% SDS polyacrylamide gels (Bio-Rad). Resolved proteins were then transferred onto nitrocellulose membranes using TransBlot Turbo System (Bio-Rad) and membranes were then blocked with 5% bovine serum albumin (BSA, Hyclone), tris-buffered saline with Tween-20 (TBST) solution. Blots were probed with primary antibodies diluted according to manufacturer’s recommended concentrations in 1% BSA/TBST overnight at 4°C with constant agitation. Blots were then washed 4 times with TBST and incubated with secondary antibodies (1:5000, 1%BSA/TBST) for 1h at room temperature. Washing steps were repeated as with primary antibodies (Supplemental Table S3) and development was achieved using Clarity Western ECL substrate (Bio-rad) incubated for 5mins with agitation at room temperature. Chemiluminescent signals were captured using Gel Doc XR+ (Bio-rad) and band intensities were analysed using Image Lab (Bio-rad) software. Uncropped scans of Western blots can be found in Supplemental Information.

### ELISA

Plasma samples were isolated and stored at -80°C after density gradient centrifugation of peripheral blood samples as mentioned under Derivation of BOECs and cell culture. Cell-free BOEC supernatant were harvested and frozen at -80°C after centrifuging at 13,000g for 10 minutes after supernatant were collected following Orbital flow setup above. Vitreous humor samples were given by Singapore National Eye Center. HYAL-1 was quantified using Human Hyaluronidase 1/HYAL1 DuoSet ELISA and DuoSet ELISA Ancillary Reagent Kit 2 (DY7358 and DY008 respectively from R&D Systems), in accordance with the manufacturer’s protocol. All plates were read by spectrophotometry at 450nm, followed by a subtraction at 540nm for optical correction using Synergy H1 (BioTek). HYAL-1 concentrations were determined from standard curves generated from four-parameter logistic (4-PL) curve fit (“Quest Graph™ Four Parameter Logistic (4PL) Curve Calculator.” AAT Bioquest, Inc) and multiplied by the dilution factor. Mann-whitney test was selected to identify significantly different HYAL-1 levels across plasma, vitreous humor and BOEC supernatant samples in both PCV and normal samples. The level of statistical significance was set at *p* value <0.05 and all statistical analysis was performed using GraphPad Prism software, version 9.0.2.

## Original Data

<u>Uncropped scans of western blots for Figure 5a</u>

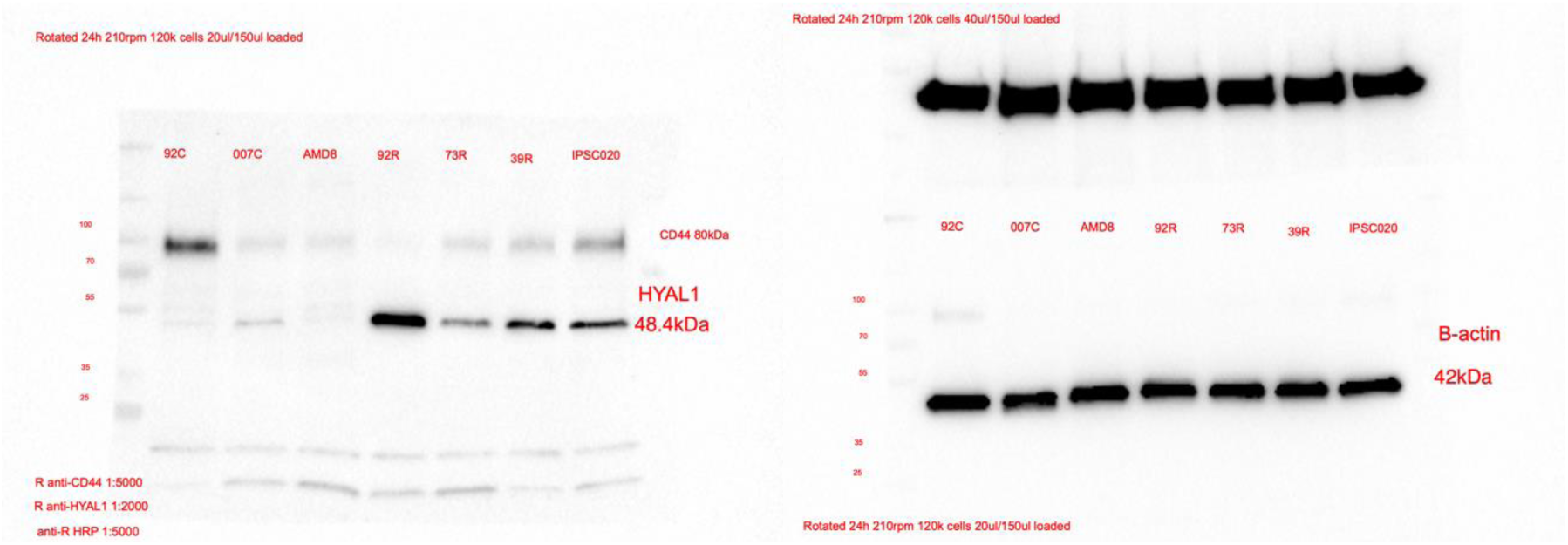

<u>Uncropped scans of western blots for Figure 6a</u>

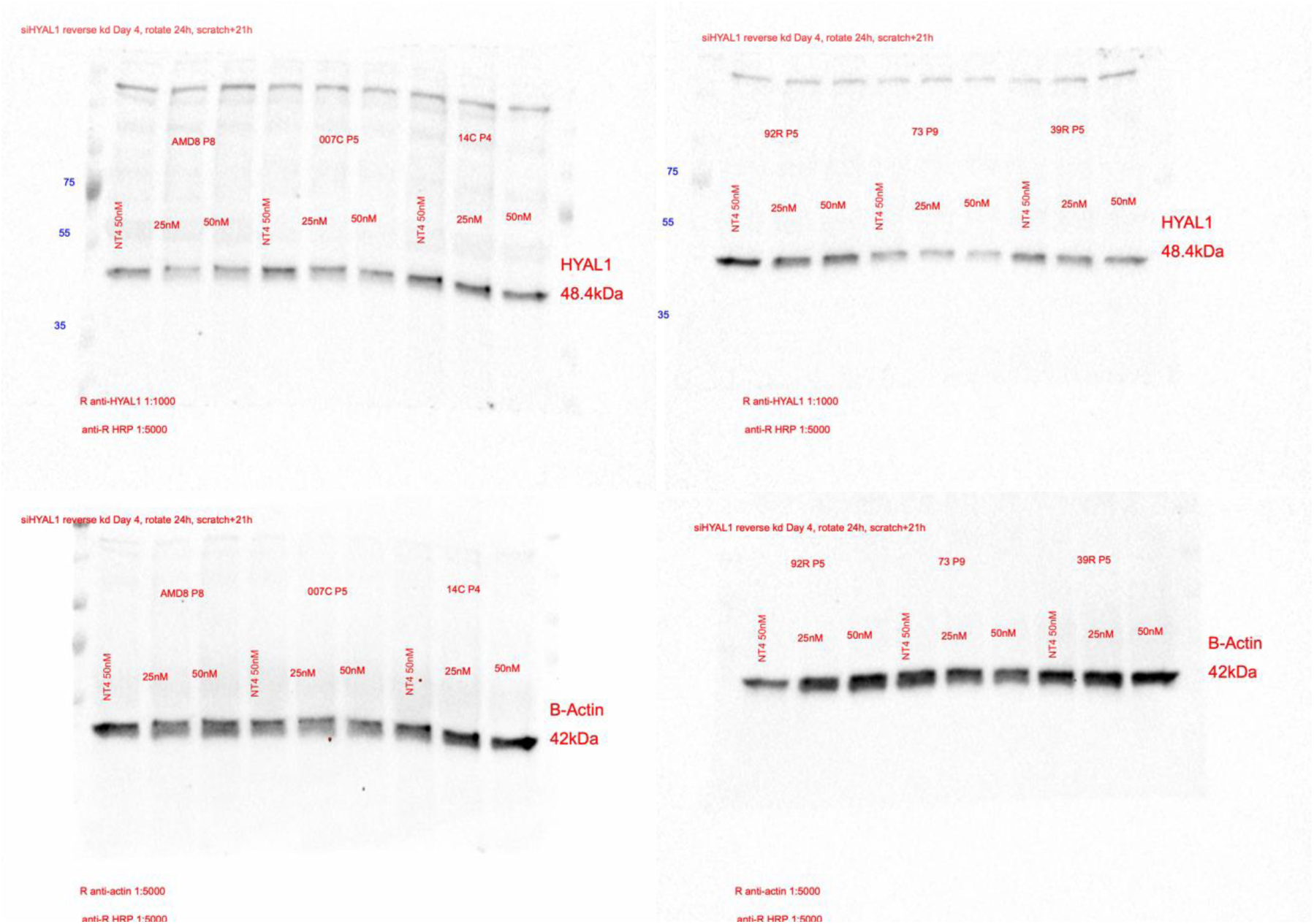

